# Mucosal immunization with DTaP confers protection against *Bordetella pertussis* infection and cough in Sprague-Dawley rats

**DOI:** 10.1101/2021.06.21.449353

**Authors:** Jesse M. Hall, Graham J. Bitzer, Megan A. DeJong, Jason Kang, Ting Y. Wong, M. Allison Wolf, Justin R Bevere, Mariette Barbier, F. Heath Damron

## Abstract

Pertussis is a respiratory disease caused by the Gram-negative pathogen, *Bordetella pertussis* (*Bp*). The transition from a whole cell pertussis vaccine (wP; DTP) to an acellular pertussis vaccine (aP; DTaP; Tdap) correlates with an increase in pertussis cases, despite widespread vaccine implementation and coverage, and it is now appreciated that the protection provided by aP rapidly wanes. To recapitulate the localized immunity observed from natural infection, mucosal vaccination with aP was explored using the coughing rat model of pertussis. Immunity induced by both oral gavage (OG) and intranasal (IN) vaccination of aP in *Bp* challenged rats over a nine-day infection was compared to intramuscular (IM)-wP and IM-aP immunized rats that were used as positive controls as IM immunization is the current route for wP and aP vaccination. Our data demonstrate that both IN and OG immunization of aP resulted in production of anti-*Bp* IgG antibody titers similar to IM-wP and IM-aP vaccinated controls post-challenge. IN-aP also induced anti-*Bp* IgA antibodies in the nasal cavity. Immunization with IM-wP, IM-aP, IN-aP, and OG-aP immunization protected against *Bp* induced cough, while OG-aP immunization did not protect against respiratory distress. Mucosal immunization (IN-aP and OG-aP) also protected against acute inflammation and decreased bacterial burden in the lung compared to mock vaccinated challenge (MVC) rats. The data presented in this study suggests that mucosal vaccination with aP can induce a mucosal immune response and provide protection against *Bp* challenge.

## INTRODUCTION

Infection of the respiratory mucosa by the Gram-negative bacterium *Bordetella pertussis* (*Bp*) causes the disease known as pertussis (whooping cough) (1). Clinical manifestations of pertussis are characterized by paroxysmal cough, hypertension, leukocytosis and in severe cases death, particularly in infants who have yet to receive their first vaccine dose (2–4). Before pertussis vaccines were introduced in the United States, pertussis led to approximately 200,000 deaths annually (5). Largely, this disease has been under control by the use of diphtheria tetanus whole-cell pertussis (DTP; wP) and acellular pertussis (DTaP; Tdap; aP) vaccines. DTP was first introduced in the 1940s/1950s, and was largely effective in decreasing pertussis incidence (6). Due to the robust immune response and reactogenicity concerns, developed countries converted to the use of DTaP in the 1990s (2).

Since the introduction of the aP vaccine, pertussis cases have been increasing, despite high vaccine coverage. It has been hypothesized that the increase in pertussis cases is attributed to: waning immunity from DTaP and Tdap vaccination, vaccine driven evolution of *Bp* strains, increased surveillance of pertussis, increased asymptomatic transmission, and improved PCR based molecular identification of cases (7–13). Infant baboons vaccinated with DTaP (1 human dose at 2, 4, and 6 months of age) were still colonized after experimental *Bp* challenge in addition to subsequent transmission to naïve baboons (14). In the same study, Warfel *et al* (2014) showed that convalescent baboons that cleared a prior *Bp* infection, were not colonized following re-challenge of *Bp* one month later (14). In humans, studies suggest that convalescent immunity can confer protection for approximately 4-20 years (15), while DTaP immunity falls short lasting on average 4-12 years (15). Overall, these data demonstrated that immunity induced through natural infection can generate longer lasting protection that also elicits pathogen clearance. As *Bp* is a respiratory pathogen, it can be hypothesized that generating an immune response at the respiratory mucosa is necessary for protection against pertussis.

Infection with *Bp* localizes to respiratory epithelium primes the immune response against subsequent *Bp* infection by recruitment of antibody producing cells and tissue resident memory T cells (Trm) (16). Bacteria invading mucosal surfaces can induce inflammation resulting in subsequent production of IgA antibodies (17). Patients previously infected with *Bp* have developed IgA antibody titers in their nasal secretions (18). In addition, anti-*Bp* IgA antibodies from patients who have convalesced from *Bp* infection inhibit bacterial attachment *in vitro* and increase *Bp* uptake and killing by human polymorphonuclear leukocytes (19, 20). Convalescent humans also generate *Bp* specific IgG antibody titers that have shown to correlate with protection (21, 22). *Bp* infected mice generate *Bp* specific CD4+ T cells in the lung that secrete IFN-γ and IL-17 (23, 24). In mice, *Bp* infection induced Trms in the lung and were associated with pathogen clearance (16). Nonhuman primates previously infected with *Bp* generate both Th1 and Th17 memory cells that are still detected two years post infection (25). To induce a similar protective immune response that is observed during natural infection, mucosal vaccination has recently been investigated (26–28)

Mucosal immunization has been scarcely used as a vaccination strategy to generate a protective immune response against pertussis in clinical and pre-clinical models. Oral vaccination with wP induced the production of *Bp* specific antibody titers in both the saliva and serum of newborns (29). In a subsequent study in 1985, oral immunization of 10^12^ CFUs of killed *Bp* led to the induction of antibody titers in the saliva and serum of newborns (30). The frequency of pertussis was lower in orally vaccinated newborns during the first year of life compared to newborns who were unvaccinated although, this difference disappeared by the end of the year (30). In mice, oral administration of attenuated bacterial vectors *Salmonella typhimurium* and *Escherichia coli* expressing *Bp* antigen, filamentous hemagglutinin (FHA), results in the production of anti-FHA IgA antibody titers in the lung (31). Intranasal immunization of a live attenuated vaccine strain of *Bp,* BPZE1 was protective both in preclinical and clinical studies (32–35). Previously, our lab has shown that IN immunization of DTaP can induce a protective immune response in mice (27, 28). Boehm *et al* (2019) illustrated that IN vaccination of DTaP with and without the addition of the adjuvant curdlan induces both anti-*Bp* and anti-pertussis toxin (PT) IgG antibody titers, as well as *Bp* specific IgA in the lung (27). A subsequent study performed by Wolf *et al* (2021) suggested that IN-aP vaccination is protective through the induction of humoral responses 6 months after booster vaccination and challenge (28). Mounting evidence supports that mucosal immunization can be protective against *Bp* colonization, but murine studies lack the ability to evaluate one the of hallmark symptoms of pertussis, *Bp*-induced coughing.

The rat model of pertussis has been utilized to characterize *Bp* pathogenesis and evaluate coughing manifestation from *Bp* infection (36–42). Only a few studies have been performed investigating vaccine efficacy in the coughing rat model of pertussis. Hall *et al* (1998) demonstrated that the SmithKline Beecham 3-component aP vaccine (detoxified PT (PTd), FHA, and a 69kDa antigen, presumably pertactin (PRN)), the Connaught 5-component vaccine (PTd, FHA, agglutinins 2+3 (fimbriae), and PRN), and Evans whole-cell pertussis vaccine protected against cough upon intrabronchial *Bp* challenge (41). Rats administered one single human dose of DTP had a lower incidence of coughing following *Bp* challenge (39). We hypothesized that both oral and IN vaccination would protect against bacterial burden upon challenge as well as protect against *Bp* induced coughing in the rat model of pertussis by generating a protective immune response at the site of infection. To test this hypothesis, we IN and OG vaccinated and challenged Sprague-Dawley rats with 1/5^th^ human dose of DTaP. IM-wP and IM-aP vaccinated and *Bp* challenged rats were used as positive controls to compare vaccine-mediated immunity as IM administration of DTaP is the current route of vaccine administration.

In our study, we aimed to use the rat model of pertussis to measure protection induced from mucosal vaccination of DTaP. By utilizing the coughing rat model of pertussis, protection against bacterial burden in the respiratory tract and prevention of *Bp* induced cough after immunization was critically analyzed. We also focused on evaluating the serological responses regarding vaccination followed by *Bp* challenge. Our data supports that protection can be afforded by mucosal immunization with DTaP. IN and OG immunization with DTaP not only induced systemic anti-*Bp* IgG antibodies but also induced mucosal anti-*Bp* IgA antibodies. These data suggest that oral vaccination of DTaP can generate a humoral immune response at the respiratory mucosa in rats. IN-aP and OG-aP was also capable of protecting against *Bp-*induced cough. Furthermore, mucosal vaccination protected against bacterial burden in the respiratory tract. In conclusion, this study highlights the benefits of using the coughing rat model of pertussis to study mucosal vaccination with *Bp* vaccines.

## RESULTS

### Intranasal immunization induces systemic anti-*Bp* IgG and anti-PT IgM and IgG antibody titers following booster immunization

We hypothesized that IN-aP and OG-aP immunization would induce systemic IgM and IgG antibodies, as mucosal immunization stimulates the induction of both systemic and mucosal antibodies (27, 28, 30). To test this hypothesis, 3-week-old Sprague Dawley rats were IM-wP, IM-aP, IN-aP, and OG-aP immunized followed by a booster vaccine at 6 weeks of age with the same corresponding vaccine (**Fig. S1**). Mucosal and systemic antibodies were measured over the course of vaccination (**Fig. S1**). Minimal differences in antibody titers prior to booster vaccination was observed for all immunized groups (**Fig. 1**). Compared to all other groups, IM-wP vaccination induced a significant increase of anti-*Bp* IgM in the serum 1-week post-booster vaccination (**Fig. 1A**). IM-wP, IM-aP, and IN-aP vaccinated rats had a significant increase in anti-*Bp* IgG antibody titers following booster immunization compared to mock vaccinated challenged (MVC) rats (**Fig. 1B**). Rats vaccinated via IM-aP had a 100-fold significant increase in anti-PT IgM antibody titers one week post booster vaccine, compared to the MVC control (**Fig. 1C**). IM-aP and IN-aP vaccinated rats had a significant increase in anti-PT IgM antibodies compared to IM-wP, and OG-aP vaccinated rats (**Fig. 1C**). This same trend was also observed in measuring anti-PT IgG antibodies (**Fig. 1D**). Here our data show that following booster immunization, IN-aP vaccinated rats developed systemic anti-*Bp* IgG and anti-PT IgG and IgM antibody titers.

**FIG 1.**
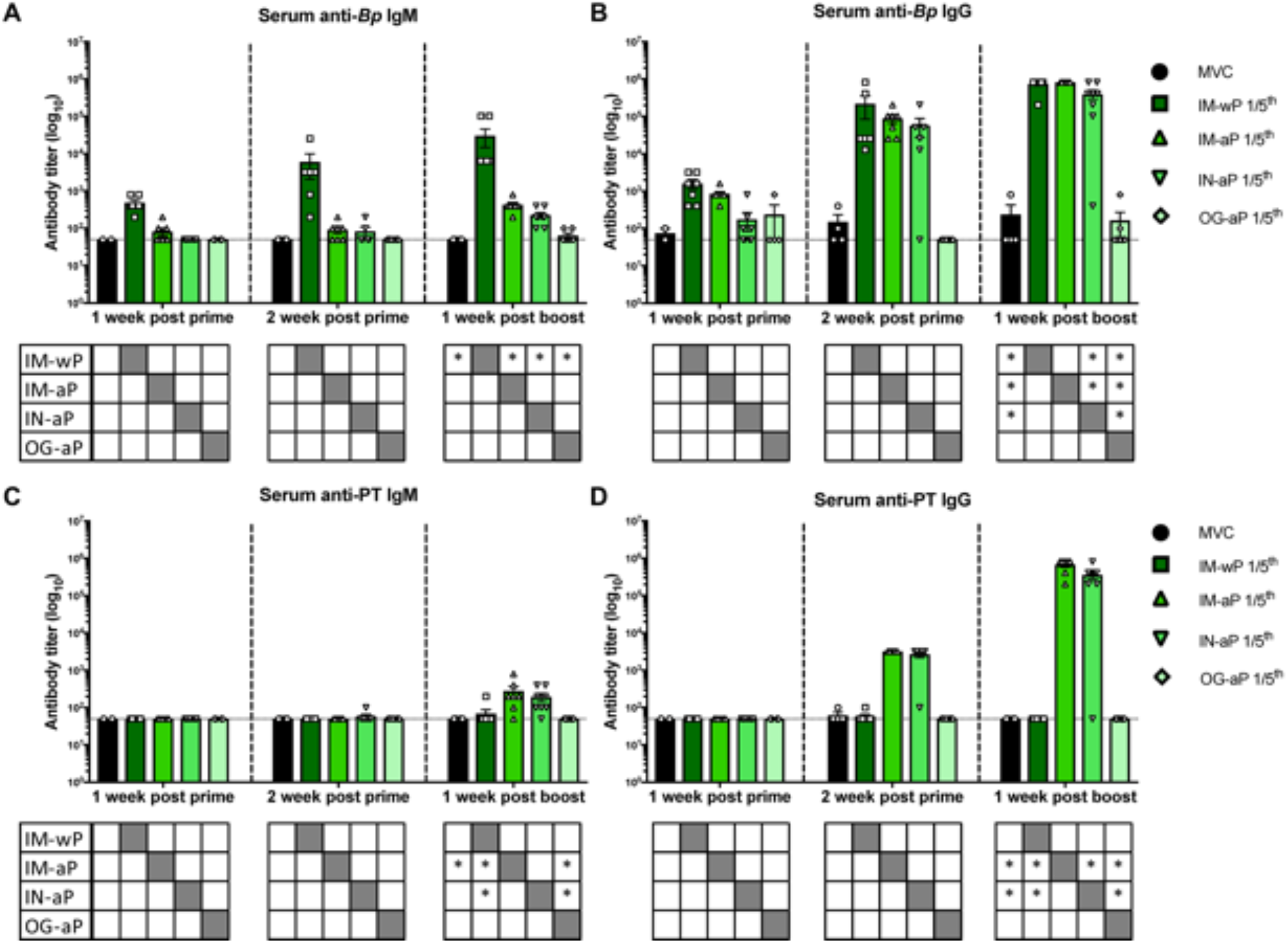
IN booster vaccination induces systemic anti-Bp and anti-PT antibody titers. 1 and 2 weeks post prime immunization and 1 week post boost blood was collected via saphenous vein, and anti Bp and anti PT IgM (A-C) and IgG (B-D) specific antibodies were measured. Results are shown on a log scale and as a mean ± SEM (n = 3-8). Dotted line represents the limit of detection. *P < 0.05. (n = 4-8). P values were determined by two-way ANOVA with Dunnett’s post hoc test compared between groups. * under each graph annotates the significance between labeled group under the y-axis and the group under the corresponding bar. Grayed out box annotates no stats calculated.

### Mucosal vaccination protects against cough from *Bp* infected rats

In 2014, Warfel *et al* showed that aP vaccination was protective against pertussis disease, but failed to protect against colonization and transmission of *Bp* in the nonhuman primate model (14). We hypothesized that mucosal vaccination with DTaP would protect against *Bp* induced cough by eliciting a protective immune response at the respiratory mucosa. To test this hypothesis, vaccinated rats were subsequently intranasally challenged with *Bp* 2-weeks post-booster vaccine administration (**Fig. S1**). Every evening post-challenge, coughs were counted using whole-body plethysmography (WBP). MVC rats averaged a total of five coughs or less during monitoring for the first 5 days of infection (**Fig. 2A**). At days 6 and 7 post-challenge, average coughs per fifteen minutes increased to more than thirty coughs (**Fig. 2A**). There was a significant decrease in coughs for rats vaccinated with IM-wP, IM-aP, IN-aP, and OG-aP at days 6 and 7 post-challenge compared to MVC rats (**Fig. 2B-E**). On average, rats vaccinated with IM-wP coughed approximately 6 coughs per fifteen minutes each day (**Fig. 2B**). Rats vaccinated with IM-aP on average coughed 2 times per fifteen minutes each day (**Fig. 2C**). IN-aP immunized rats on average coughed 3 times per fifteen minutes each day, while OG-aP vaccinated rats coughed on average 4 times per fifteen minutes each day (**Fig. 2D&E**). To compare the average total number of coughs each day post-challenge, we calculated the total number of coughs for each group per animal. We observed a significant decrease in total number of coughs in rats vaccinated with IM-aP (14 coughs) and IN-aP (27 coughs) compared to MVC rats (102 coughs) over the nine-day infection (**Fig. 2F**). OG-aP immunized rats coughed on average 35 times (**Fig. 2F**). Our data demonstrates that mucosal vaccination in rats protects against *Bp* induced cough.

**FIG 2.**
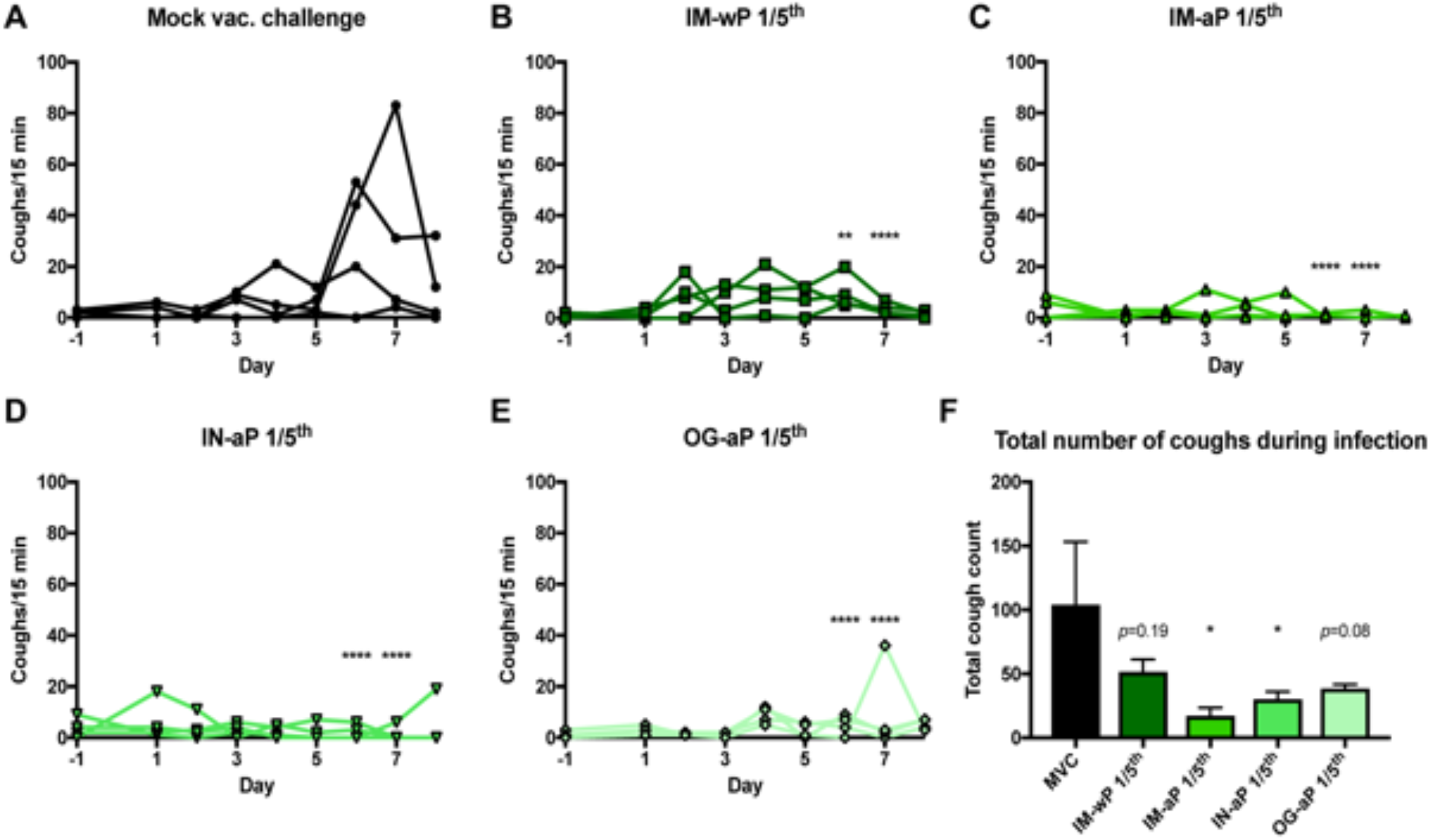
Intranasal and oral vaccination of acellular pertussis vaccine decreases cough of *B. pertussis* infected rats. Coughs were counted every day of the nine-day infection using whole body plethysmography. Coughs were counted for (A) mock vac. challenge rats, (B) IM-wP (C) IM-aP (D) IN-aP, and (E) OG-aP, vaccinated and challenged rats. To assess any potential differences between vaccine groups over the entire course of infection, (F) average total number of coughs for each rat per group was compared. Results shown as mean ± SEM (*n* = 3-4). *P* values were determined by two-way ANOVA with Dunnett’s post hoc test and one-way ANOVA with Dunnett post hoc test for total cough count, \**P* < 0.05, \*\**P* < 0.01, \*\*\**P* < 0.001, \*\*\*\**P* < 0.0001 compared to the mock vac. challenged control group.

### Intranasal vaccination protects against pulmonary distress

Our previous work has shown that *Bp* infected rats had a significant increase in pulmonary distress following challenge (43). Pulmonary distress can be evaluated by calculating enhanced pause (PenH). PenH functions as a representation of bronchoconstriction taking into consideration the timing between early and late expiration and the estimated maximum inspiratory and expiratory flow per breath. We hypothesized that mucosal vaccination would protect against *Bp* induced pulmonary distress, as bacterial clearance would decrease inflammation. Here, rats vaccinated with IM-wP, IM-aP, and IN-aP had a significant decrease in PenH compared to the MVC control group at days 5 and 7 post-challenge (**Fig. 3A-D**). Rats vaccinated IM-aP and IN-aP also had a significant decrease in PenH at day 6 post-challenge compared to MVC rats (**Fig. 3A&C-D**). However, there was no significant decrease in PenH in OG-aP vaccinated rats compared to MVC suggesting that the induced immune response is not sufficient enough to protect rats from *Bp* induced respiratory distress (**Fig. 3E**). Other respiratory parameters were also measured using WBP (**Fig. S2**). In brief, we observed that rats vaccinated IM-wP, IM-aP, and IN-aP had a significant decrease in pause (PAU) compared to MVC control group, which is another indicator of bronchiole restriction (**Fig. S2B**). Rats vaccinated with the aP regardless of route also had a significant decrease in Tidal volume (TVb) compared to MVC rats, which could be crudely associated with inflammation (**Fig. S2D**). Overall, our data demonstrated that IN-aP vaccination decreases pulmonary distress of *Bp* infected rats.

**FIG 3.**
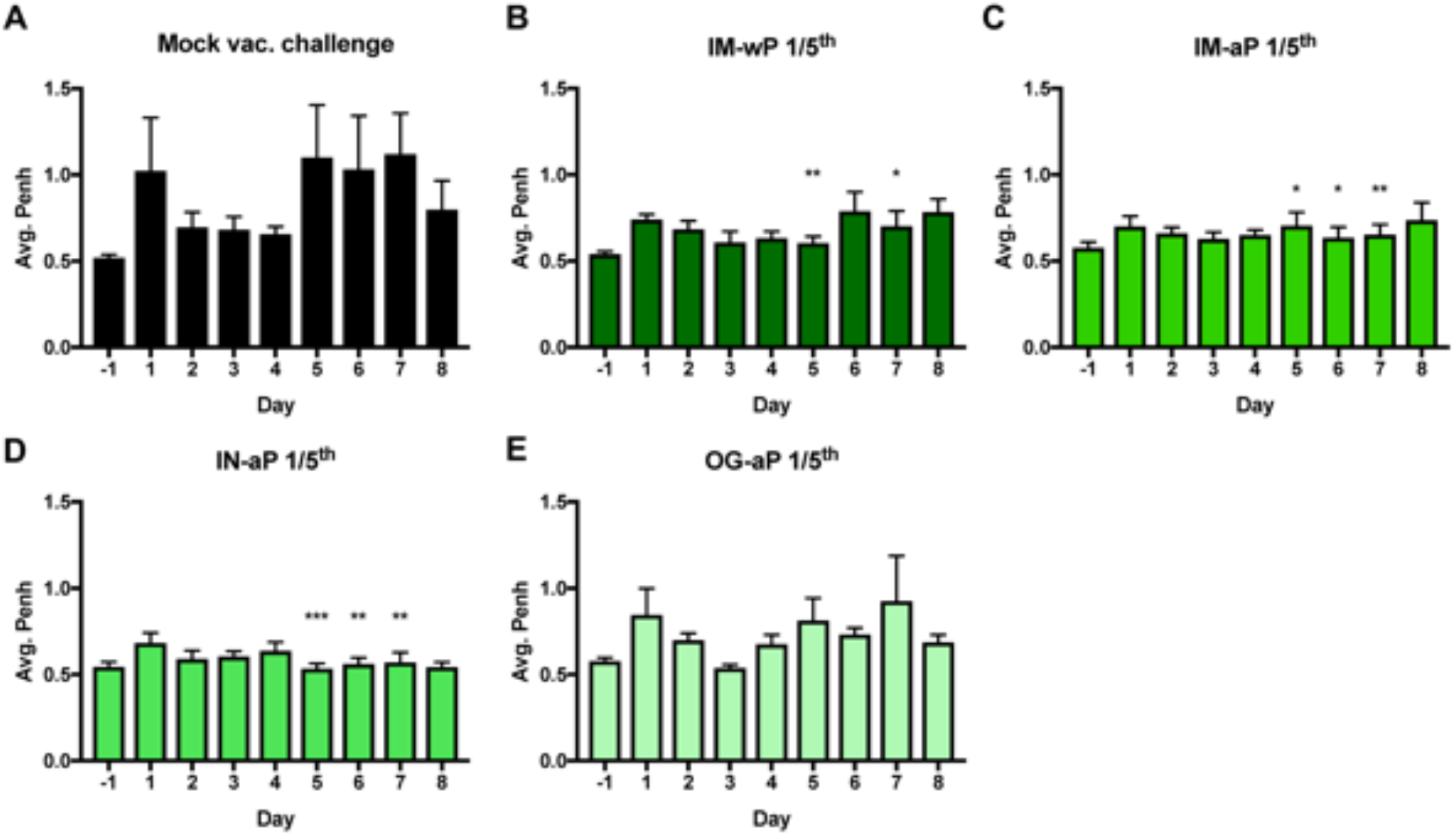
Intranasal vaccination decreases pulmonary restriction of *Bordetella pertussis* infected rats. Bronchiole restriction was measured over the course of infection by whole body plethysmography. Bronchiole restriction was determined by the factor Penh for (A) mock vac. challenge rats, (B) IM-wP (C) IM-aP (D) IN-aP, and (E) OG-aP vaccinated and challenged rats. Results shown as mean ± SEM (*n* = 3-4). *P* values were determined by two-way ANOVA with Dunnett’s post hoc test, \**P* < 0.05, \*\**P* < 0.01, \*\*\**P* <0.001 compared to the mock vac. challenge group.

### Mucosal immunization induces production of *Bp* specific antibodies in the serum, while intranasal immunization also induces PT specific antibodies in the serum

Next, we wanted to measure systemic antibody responses to *Bp* and PT following challenge. IM-wP vaccinated rats had a slight increase in anti-*Bp* IgM antibodies compared to all other vaccinated groups, albeit not significant (**Fig. 4A**). We observed a significant increase in anti-*Bp* IgG antibody titers in IM-wP, IM-aP, and IN-aP vaccinated rats compared to the MVC at day 1 post challenge (**Fig. 4B**). At day 9 post-challenge, all vaccinated rats had a significant increase of anti-*Bp* IgG antibody titers compared to the MVC control (**Fig. 4B**). Following *Bp* challenge, IM-aP and IN-aP immunized rats had a significant increase in anti-PT IgM antibodies compared to IM-wP immunized rats and MVC control. (**Fig. 4C**). Similar results were observed in measuring anti-PT IgG titers (**Fig. 4D**). IM-aP and IN-aP vaccination induced a significant increase in anti-PT IgG antibody titers compared to MVC, IM-wP, and OG-aP immunized rats after booster vaccination and at days 1 and 9 post-challenge (**Fig. 4D**). Although not significant, two of the OG-aP immunized rats had detectable anti-PT IgG antibody titers in the serum at day 9 post-challenge (**Fig. 4D**). Enzyme-linked immune absorbent spot (ELISpot) assay was used to determine the number of *Bp* specific IgG cells in the bone marrow at day 9 post-challenge. There was an increase in the number of *Bp* specific IgG cells in the bone marrow in all vaccination groups compared to the MVC control; however, the only significant increase in *Bp* specific IgG producing cells in the bone marrow was detected in IM-wP vaccinated rats (**Fig. S3**). Our data indicate that mucosal vaccination via IN and OG immunization induced *Bp* specific IgG antibody responses.

**FIG 4.**
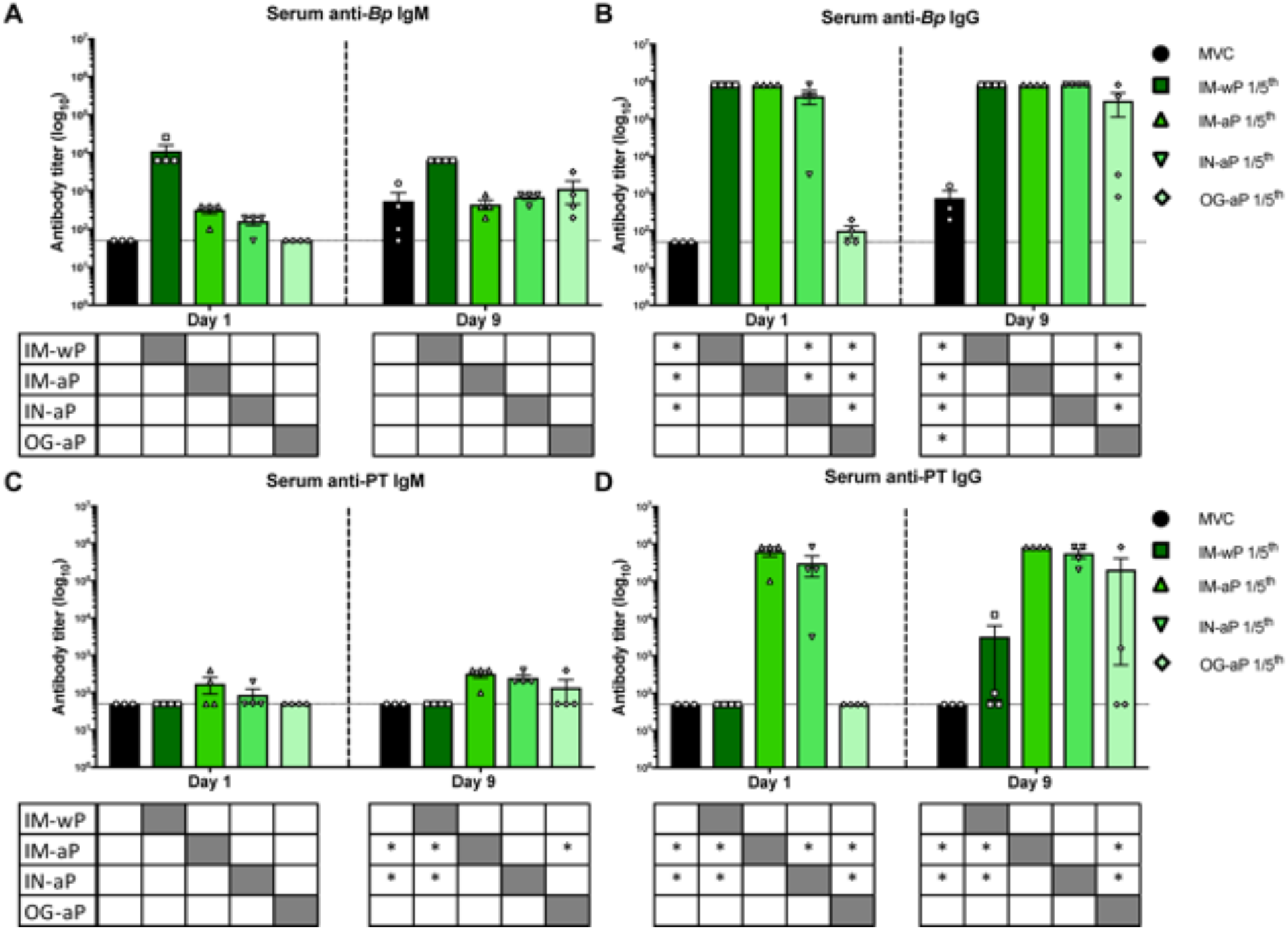
Mucosal vaccination induces production of anti-*Bp* IgG, while IN immunization also induces both anti-PT IgM and IgG antibodies. ELISAs were used to determine and compare the induced serological responses from vaccinated and challenge rats in the serum. Both (A, C) IgM and (B, D) IgG serum antibody titers from immunized and challenged rats were measured post prime, boost, and challenge. Dotted line represents the limit of detection. Results are shown on a log scale and as mean ± SEM, \**P*< 0.05 (*n* = 4). P values were determined by two-way ANOVA with Dunnett’s post hoc test compared between groups. * under each graph annotates the significance between labeled group under the y-axis and the group under the corresponding bar. Grayed out box annotates no stats calculated.

### Intranasal immunization induces production of *Bp* specific IgA antibodies in the nasal cavity

In humans, previous *Bp* infection leads to anti-*Bp* IgA antibodies in nasal secretions (18). IgA antibodies to *Bp* play a role in the inhibition of *Bp* attachment *in vitro* to epithelial cells (19). Here, we investigated if IN and OG immunization of DTaP would induce mucosal IgA antibodies in the lung and/or the nasal cavity. In the lung, three of the four IN-aP vaccinated rats had detectable anti-*Bp* IgA antibodies at day 1 post-challenge, although not significant (**Fig. 5A**). We did not detect anti-*Bp* IgA antibody titers at day 1 post-challenge in the lung of IM-wP, IM-aP, or OG-aP vaccinated rats (**Fig. 5A**). Low levels of anti-*Bp* IgA titers were measured in all vaccinated groups at day 9 post-challenge albeit not significant compared to our MVC control (**Fig. 5A**). The same trend was observed in the lung measuring anti-PT IgA titers at day 1 post-challenge (**Fig. 5B**). At day 9 post-challenge 50% of IN-aP rats and 25% of rats OG-aP vaccinated had detectable anti-PT IgA (**Fig. 5B**). In the nasal cavity, only one IN-aP vaccinated rat had detectable anti-*Bp* IgA antibody titers; however, we did measure a significant increase in anti-*Bp* IgA antibodies in IN-aP immunized rats at day 9 post-challenge compared to the MVC, IM-wP, IM-aP, and OG-aP immunized rats (**Fig. 5C**). Only one IN-aP and one OG-aP vaccinated rat had detectable amounts of anti-PT IgA in the nasal cavity at day 9 post challenge (**Fig. 5D**). Overall, our data reveal that IN-aP immunization is capable of inducing IgA antibodies in the nasal cavity of rats.

**FIG 5.**
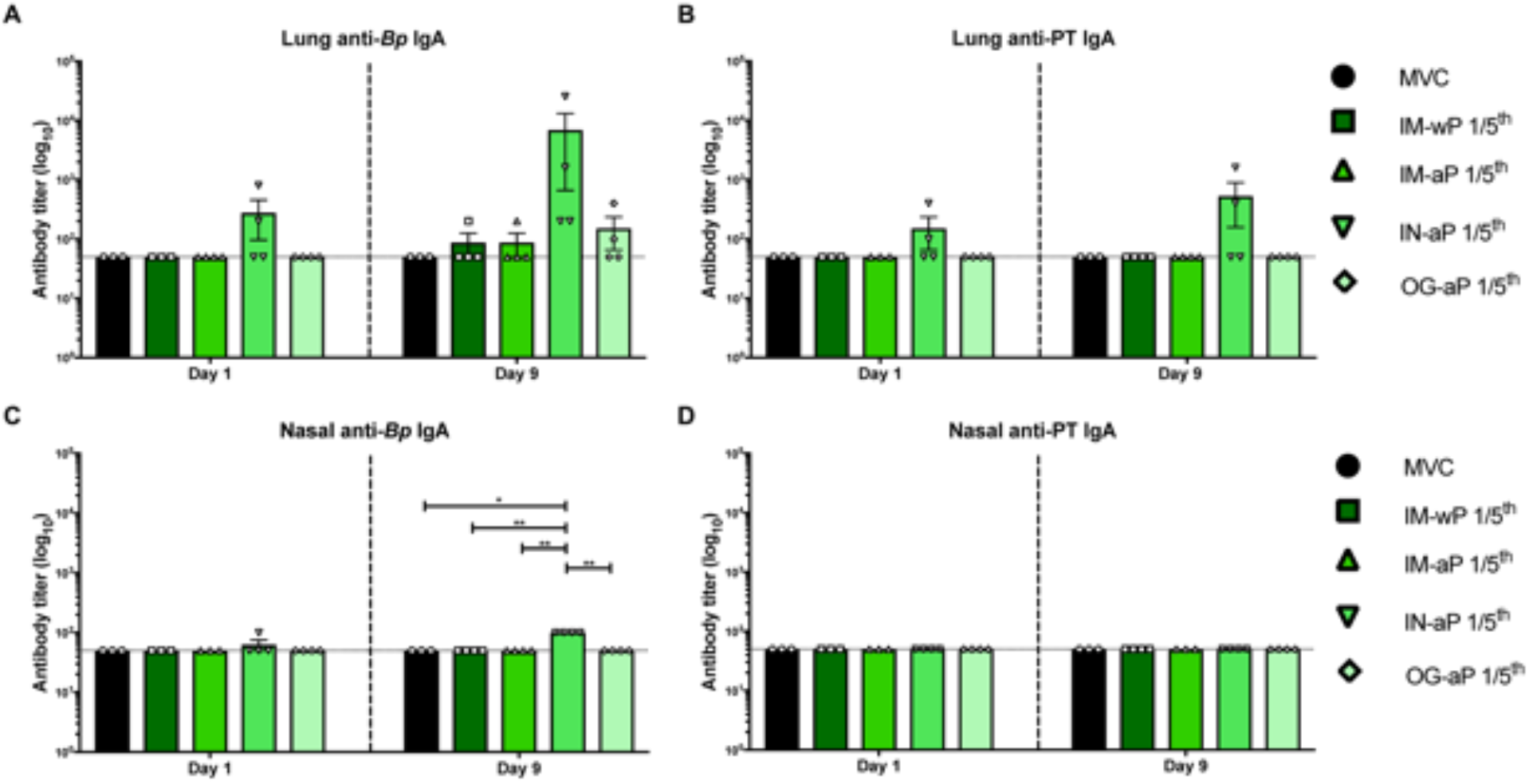
Intranasal immunization elicits the production of anti-*Bp* IgA in the respiratory tract. ELISA was used analyze antibodies in the (A, B) lung and (C, D) nasal cavity from lung homogenate supernatant and PBS flushed through the nasal cavity from vaccinated and challenge rats at days 1 and 9 post challenge. IgA titers were determined against pertussis toxin and *B. pertussis*. Dotted line represents the limit of detection. Results are shown on a log scale and as a mean ± SEM (n = 3-4). **P < 0.01, ****P < 0.0001 *P* values were determined by Kruskal-Wallis test with Dunnett’s post hoc test compared between groups.

### Mucosal immunization protects against acute inflammation in the lung

Our previous rat challenge study illustrated that intranasal *Bp* challenge gave rise to both acute and chronic inflammation in the rat lung (43). Here, we used histology to assess if mucosal immunization would protect against *Bp* induced inflammation in the lung. At day 1 post-challenge, no differences in acute inflammation were observed; however, at day 9 post-challenge, rats vaccinated with IM-aP, IN-aP, and OG-aP had a significant lower acute inflammation scoring compared to the MVC rats. (**Fig. 6A&C**). IN-aP immunized rats had a higher chronic inflammatory score at day 1 post-challenge (**Fig. 6B&D**). There were no observed differences in chronic inflammation in vaccinated rats compared to MVC rats at day 9 post-challenge (**Fig. 6D**). Total inflammation was calculated by combining both acute and chronic inflammatory scores, as rat lungs exhibited both types of inflammation. No differences in total inflammation score were observed at day 1 post-challenge; however, all vaccinated rats had a significant lower total inflammation score compared to MVC rats (**Fig. 6E**). There were no differences in lung weight, which can be used as a crude measurement for lung inflammation, following *Bp* challenge. (**Fig. S4A**). We did observe a significant increase in percent body weight change in IM-wP vaccinated rats compared to MVC control rats suggesting that wP protects against weight loss observed in non-vaccinated challenged rats (**Fig. S4B**). Our observations suggest that that mucosal vaccination protects against *Bp* induced inflammation in the lung (**Fig. 6C&E**).

**FIG 6.**
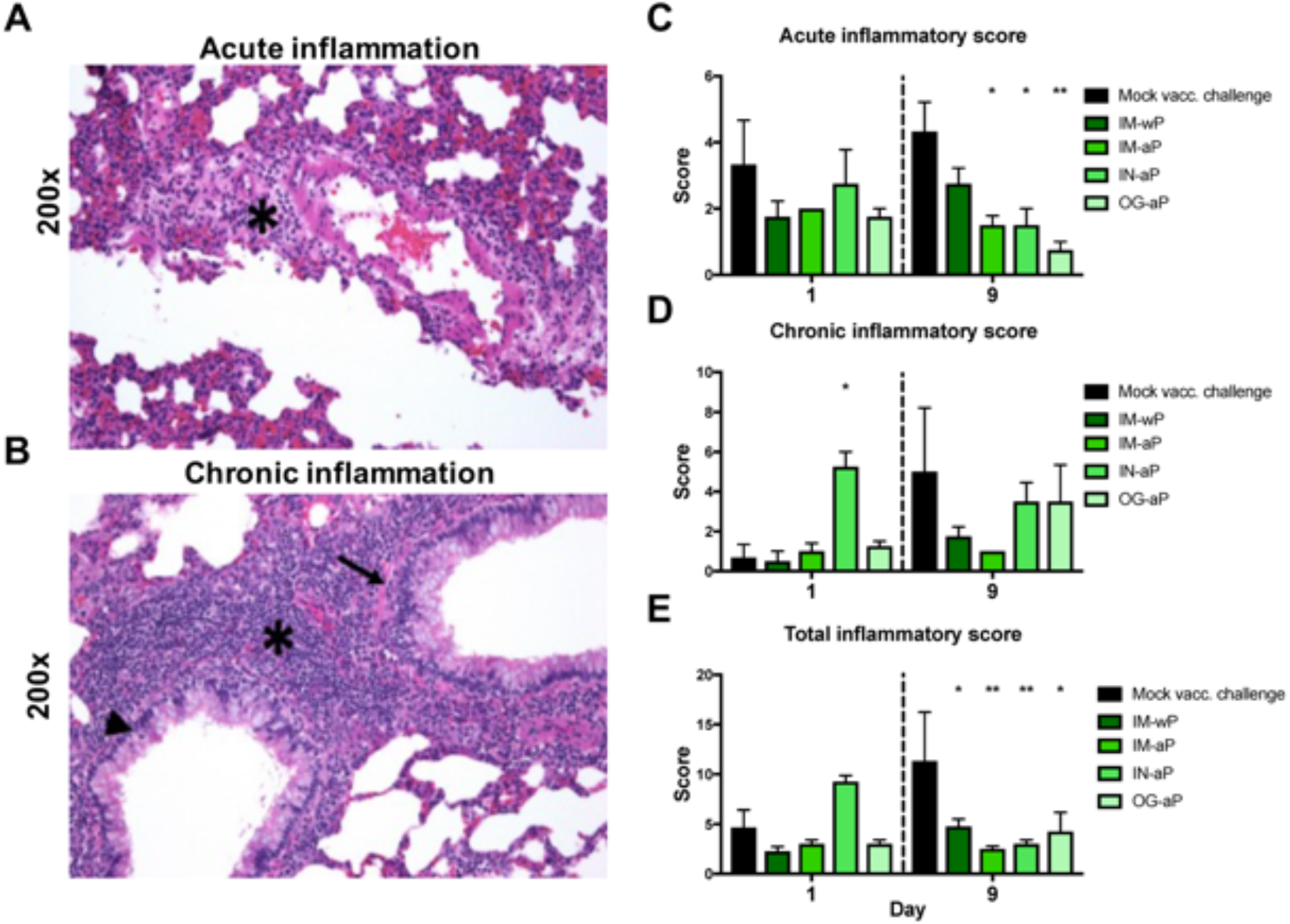
Mucosal vaccination protects against acute and total inflammation in the lung of *Bp* infected Sprague-Dawley rats. Post euthanasia, left lobe of the lung was excised, sectioned, and stained with hematoxylin and eosin. Lung samples scores were based on standard qualitative toxicologic scoring criteria (0 – none, 1 – minimal (rare), 2 – mild (slight), 3 – moderate, 4 – marked, 5 – severe). (A) Representative image of acute inflammation of rat lung showing increased numbers of neutrophils and edema surrounding blood vessel (asterisk). (B) Representative image of chronic inflammation of the rat lung showing increased numbers of mononuclear cells surrounding bronchioles (asterisk). Inflammatory cells are also present in the lamina propria (arrow) and epithelium (arrowhead) of bronchioles. (C) Average acute inflammation scores of the lung are detailed by the presence of neutrophils in the parenchyma, blood vessels, and the airways. (D) Average chronic inflammation scores are distinguished by mononuclear infiltrates in the parenchyma, blood vessels, and airway of the lung. (E) Total inflammatory score calculated by the sum of the acute and chronic inflammation score of the lung. All scoring assessments were determined with no knowledge of the groups. Results are shown as mean ± SEM (*n* = 3-4) *P* values were determined by two-way ANOVA followed by Dunnett’s comparison test, \**P* < 0.05, \*\**P* < 0.01 compared to mock challenge.

### Mucosal vaccination protects against *Bp* challenge

Next, we wanted to assess if mucosal immunization could protect against *Bp* burden in the respiratory tract. Bacterial burden in the respiratory tract was determined 1hr, 1-, and 9- days post *Bp* challenge. Bacterial burden was measured at 1hr post-challenge (n=2) to assess potential bacterial loss for our original challenge dose. In the lung, trachea, and nasal lavage fluid, we measured approximately 10^6^ CFUs 1hr post challenge (**Fig. 7A-C**). At day 1 post-challenge there was a significant 98.5% reduction in bacterial burden in the lung of IM-aP immunized rats compared to MVC (**Fig. 7A**). IM-wP. IM-aP, IN-aP, and OG-aP vaccinated rats all had a significant decrease in bacterial burden in the lung at day 9 post-challenge compared to MVC rats (**Fig. 7A**). There was also a significant decrease in bacterial burden in the trachea at both days 1 and 9 post-challenge in all vaccinated rats compared to MVC (**Fig. 7B**). At day 1 post-challenge, there was a significant 86-97% reduction in bacterial burden in all vaccinated rats compared to MVC rats in the nasal cavity (**Fig. 7C**). At day 9 post-challenge we did not measure any significant differences between groups as most of the bacteria were cleared from the nasal cavity (**Fig. 7C**). Overall, we observed that IN-aP and OG-aP vaccinated rats have a significant reduction in bacterial burden in the respiratory tract compared to the MVC control group at days 1 and 9 post-challenge (**Fig. 7A-C**).

**FIG 7.**
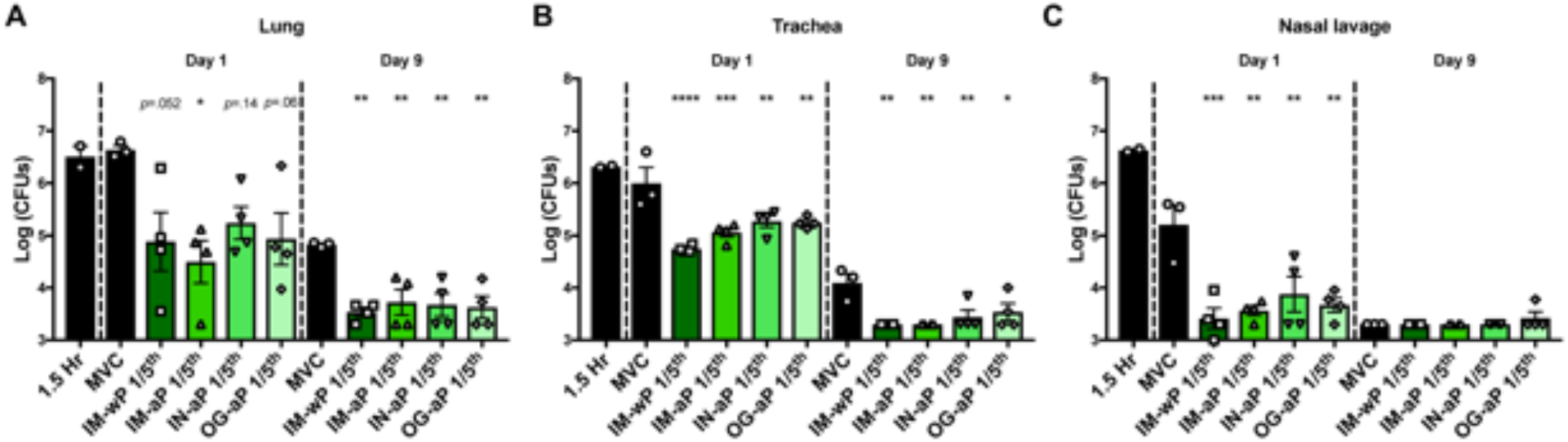
Oral and intranasal immunization decreased *B. pertussis* bacterial burden in the respiratory tract. Bacteria were quantified by serially diluted CFUs following vaccination and intranasal challenge. CFU counts were determined from (A) lung homogenate (B) trachea and (C) nasal lavage 1.5Hr, 1-, and 9-day post *B. pertussis* challenge. Results are shown as mean ± SEM (*n* = 2-4). *P* values were determined by one-way ANOVA with Dunnett’s post hoc test, \**P* < 0.05, \*\**P* < 0.01, \*\*\**P*<0.001, \*\*\*\**P* < 0.0001 compared to mock vac. challenge group.

### wP immunization induces a proinflammatory cytokine response compared to mucosal vaccinated Sprague-Dawley rats

Previous studies have shown that both wP immunization and *Bp* infection induces a pro-inflammatory Th1/Th17 immune response, while aP immunization promotes a more Th2 skewed response (44–50). In our current study, we measured cytokines in the lung and serum induced from vaccination and challenge. In the lung at day 1 post-challenge, we measured a significant 4-fold increase in IL-17 in MVC rats compared to IM-aP and IN-aP vaccinated rats (**Fig. 8A**). At day 9 post-challenge, IM-wP immunized rats had a significant increase in IL-17 compared to IM-aP, IN-aP, and MVC rats (**Fig. 8A**). IM-wP vaccinated rats also had a significant increase in Th1 cytokine IL-12p70 compared to MVC rats in the lung and serum at day 9 post-challenge (**Fig. 8A&B**). IM-wP vaccinated rats had a significant increase in Th2 cytokines IL-4 and IL-13 in the serum compared to IM-aP and IN-aP vaccinated rats and a significant increase in IL-4 to MVC control at day 9 post-challenge (**Fig. 8B**). IM-wP immunized rats also had a significant increase in G-CSF in the serum day 9 post-challenge compared to IM-aP, IN-aP, OG-aP, and MVC rats. Overall, we did not observe marked changes in cytokine responses between DTaP vaccinated rats compared to the MVC control group; however, rats OG-aP immunized did have a slight increase in IL-17, though not significant. The observed difference in cytokines levels could be from *Bp* challenge rather than vaccination. The increase in acute inflammatory score in the lung of IM-wP immunized rats at day 9 post-challenge could be associated to the increase in proinflammatory cytokines. Graphs showing the statistical significance between cytokines in the serum and lung are in the supplementary data (**Fig. S5&6**). Our data support that response to *B. pertussis* in IM-wP immunized animals is associated robust cytokine response compared to both naïve and aP vaccinated rats, which is to be expected based on the work that has examined the Th17 response induced by whole cell pertussis vaccines.

**FIG 8.**
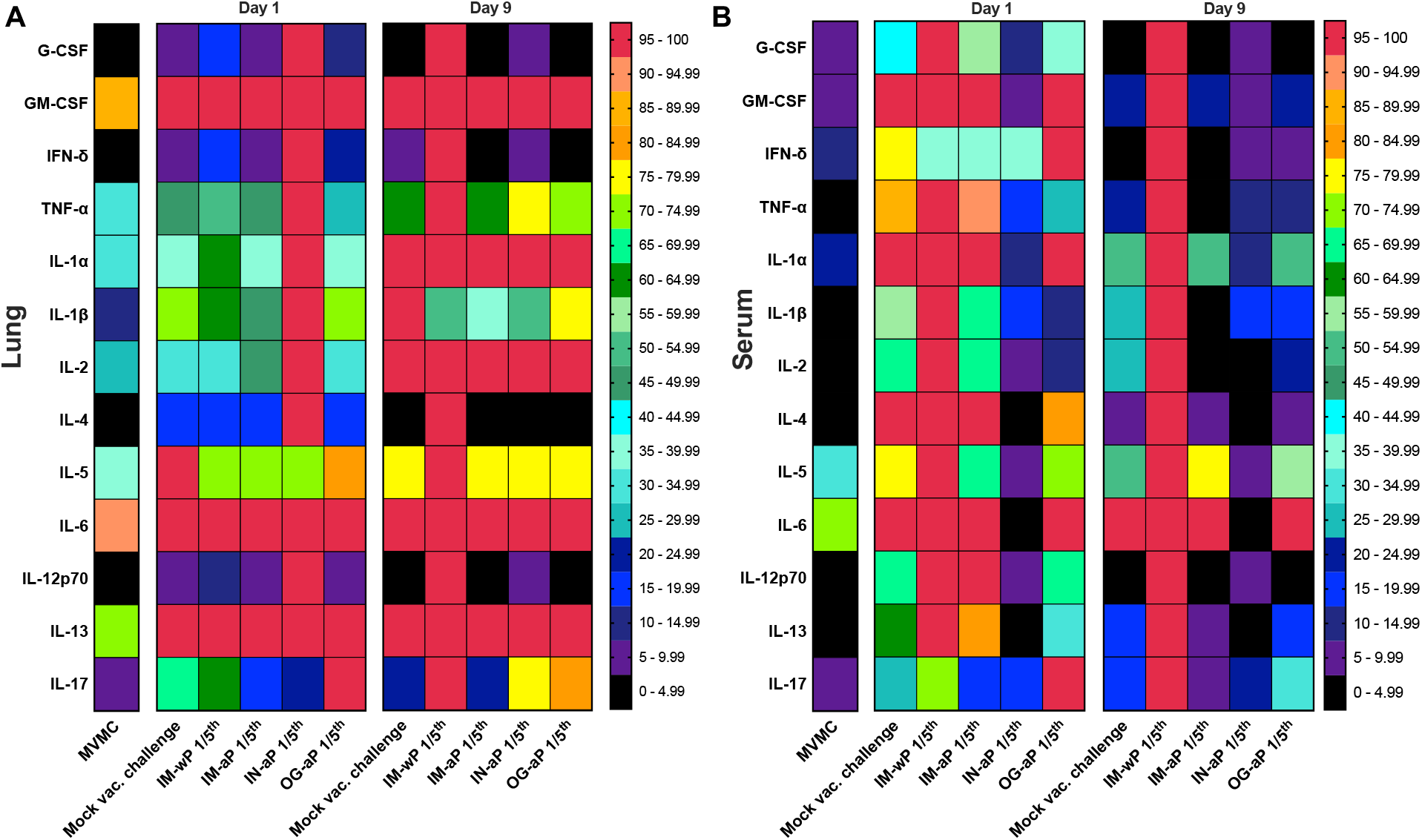
Measurement of cytokines in the lung and serum at days 1 and 9 post infection. Heat map of the average percent cytokines normalized to the max cytokine measured in the (A) lung and (B) serum. MVMC (mock vaccinated mock challenge) cytokines are from rats in (Hall et al 2021). All statistical analysis comparing average cytokine values are in Fig S5-6.

*Bp* infection induces an increase in circulating neutrophils in the blood, as well as white blood cells and lymphocytes (51–56). In our current study, we utilized hematology and flow cytometry to evaluate these populations. Hematology analysis revealed a significant increase in blood lymphocytes in the IN aP vaccinated rats compared to MVC post-challenge; however, no other differences in white blood cell counts in the blood were observed in the other vaccinated groups. (**Fig. S7A&B**). At day 1 post-challenge, there was a significant decrease in circulating neutrophils in the blood for IN-aP immunized rats compared to MVC rats (**Fig. S7C**). Flow cytometry analysis observed minimal differences in the number of neutrophils and B cells at days 1 and 9 post-challenge in all groups (**Fig. S7E&F**). Based upon these data, subtle differences in various circulating cell populations were observed following IN-aP vaccination.

### Serological responses correlate with bacterial clearance in the respiratory tract

Currently no definitive correlates of protection (CoP) for vaccines to protect against *Bp* have been established (22). It is appreciated that Th17 responses as well as Trms correlate with strong protection in mice and baboons. Antibodies to PT/FHA/PRN do not always correlate with protection in humans. In an effort to more precisely define correlates, using the rat model, we aimed to utilize the coughing phenotype and bacterial burden to identify the nature of how each vaccine protects (OG/IN/IM; acellular or wP). Previous work in our lab performed by Wolf *et al* (2021) illustrated that serum anti-*Bp*, anti-FHA, and anti-PT IgG antibody titers in the serum following IN vaccination in mice correlate with the decrease in bacterial burden in the lung following *Bp* challenge (28). Previous studies have shown that serum anti-*Bp* IgG antibodies induced from wP vaccination correlate with protection against bacterial burden in the lung of *Bp* challenged mice (57). Here, we hypothesized that antigen specific serum IgG and mucosal IgA antibodies correlate with decreased bacterial burden and cough, as IN-aP and OG-aP vaccination induced systemic and mucosal antibody responses. To test this hypothesis, we generated correlograms to evaluate both negative and positive correlations elicited by each vaccination route (58). Correlograms are an analysis tool that can be used to determine if the relationship observed between variables (i.e. bacterial burden and antibody titers) is random or not (59). If the relationship between the two variables is random the R^2^ correlation value is or near zero (59). The relationship is considered correlative if the R^2^ values approximately positive or negative one (59). Significant positive nonzero correlation values demonstrate a positive correlation between variables, while significant negative nonzero values represent a negative correlation (59). Negative correlations are observed when two variables are inversely related to one another; that is, when one variable increases, the other decreases. With these data, as bacterial burden would drop, then the correlate would increase (negative correlation; inverse). A positive correlation would mean that as bacterial burden increases so does the correlate that is being compared to.

IM-wP vaccinated rats had strong negative correlations (protective) between serum anti-*Bp* IgG antibodies to both bacterial burden in the lung (R^2^= −0.97) at day 1 post challenge and the nasal cavity (R^2^ = −0.84) at day 9 post-challenge (**Fig. 9A-B**). At day 1 post-challenge, we observed negative correlations (protective) between systemic anti-*Bp* IgM and anti-PT IgG antibodies in IM-aP vaccinated rats to bacterial burden in the nasal cavity (R^2^ = −0.64, −0.64 respectively). Additionally, negative correlations were observed between anti-*Bp* IgM antibodies and bacterial burden in the trachea (R^2^ = −0.72) at day 9 post-challenge (**Fig. 9C-D**). IM-aP vaccinated rats also had negative correlation between total cough counts over the course of challenge with IgG and IgM antibodies to *Bp* and PT (**Fig. 9D**). We expected that IN immunization would induce negative correlations between both serum- and mucosal-specific antibodies compared to bacterial burden in the lung, trachea, and nasal cavity at day 1 post-challenge (**Fig. 9E**). Lung anti-*Bp* IgA antibodies also negatively correlated with total cough count (R^2^ = −0.74) and bacterial burden (R^2^ = −0.6) in the lung at day 9 post-challenge for IN-aP vaccinated rats (**Fig. 9F**). We observed strong negative correlations between serum IgG and mucosal IgA antibody to bacterial burden in the lung (R^2^ = −0.93, −0.93 respectively) in OG-aP vaccinated rats at day 1 post-challenge despite the overall lower serological responses (**Fig. 9G**). At day 9 post challenge, there was a negative correlation between serum anti-*Bp* IgG antibodies to bacterial burden in the lung (R^2^ = −0.81), trachea (R^2^ = −0.49), and nasal cavity (R^2^ = −0.52) in OG-aP immunized rats (**Fig. 9H**). We also noticed that OG-aP vaccinated rats had negative correlations between serum IgG and mucosal IgA antibodies in the lung to total cough count (R^2^ = −0.94, −0.84) at day 9 post-challenge (**Fig. 9H**). Positive correlations (non-protective) were observed between bacterial burden and total inflammatory score in IM-wP, IM-aP, and OG-aP immunized rats (**Fig. 9A, C, G**). Bacterial burden also positively correlated with total cough counts at day 9 post challenge (**Fig. 9B, D, F, H**). By utilizing correlograms, strong negative correlations between serum serological responses and bacterial burden were observed in IM-wP and IM-aP immunized rats. Additionally, our data underscore the idea that both systemic and mucosal antibodies correlate with the observed *Bp* clearance in the respiratory tract and protection from *Bp* induced cough elicited from IN-aP and OG-aP vaccination highlighting the observed differences between vaccination routes.

**Fig 9.**
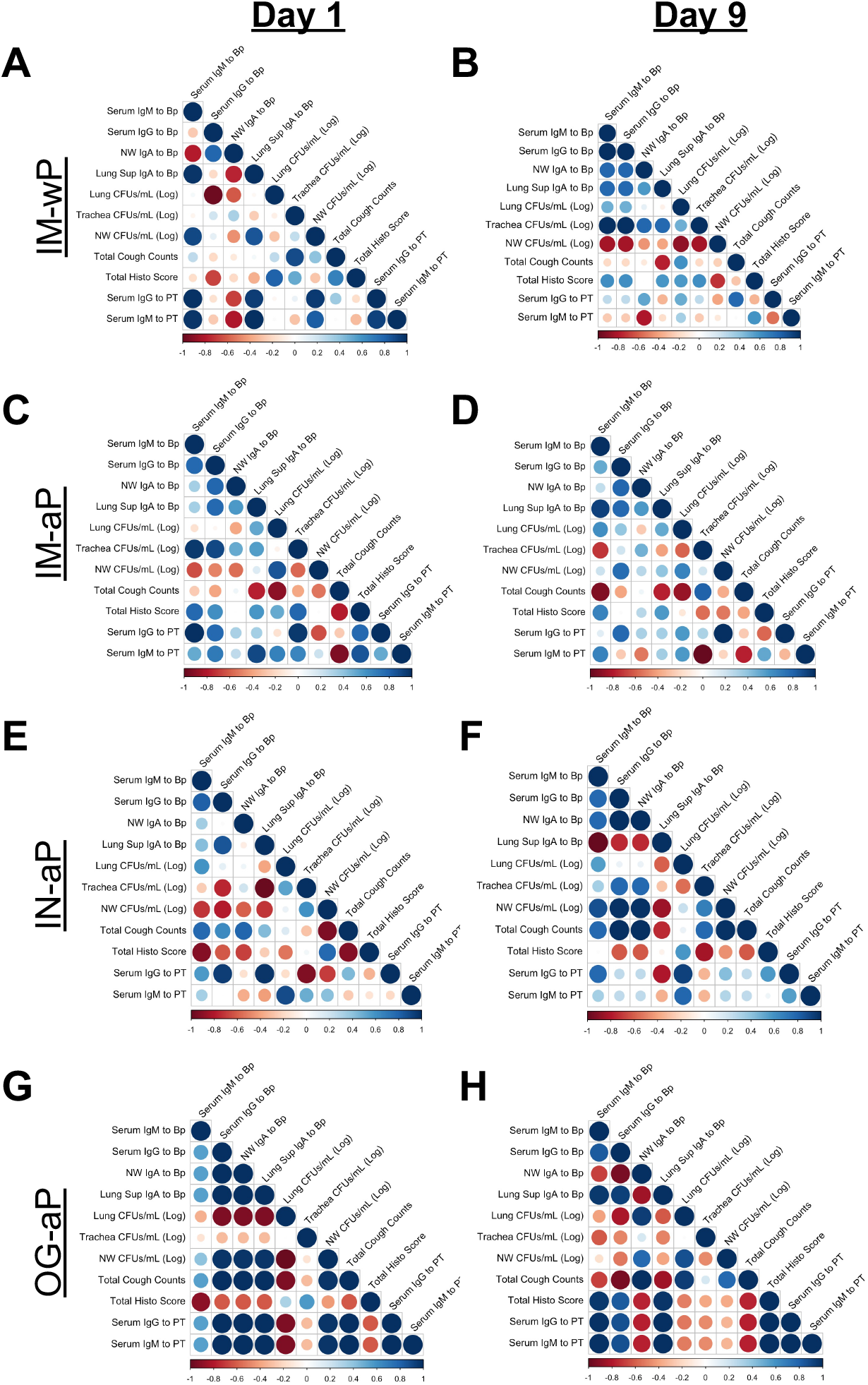
Systemic and mucosal anti-*Bp* and anti-PT antibodies correlate with observed protection. Correlograms were generated using the observed data for IM-wP (A-B), IM-aP (C-D), IN-aP (E-F), and OG-aP (G-H). Program R was used to make correlation graphs from raw data for both day 1 and day 9 post-challenge. R^2^ values were generated when generating the correlograms. Positive correlations are annotated by the blue circles, while the negative correlations are annotated by the red circles. The size of the circle annotates the strength of the correlation.

## Discussion

The immunity induced by aP vaccines is relatively short lived; thus, DTaP/Tdap vaccinated individuals are still capable of *Bp* transmission (60, 61). We have recently re-investigated the rat model of pertussis to further understand *Bp* pathogenesis from current circulating *Bp* strains (such as CDC isolate D420) (43). The coughing rat model of pertussis is a tool that can be used to evaluate bacterial burden in the respiratory tract, and also evaluate vaccine-induced immunity against cough and respiratory function (36–41, 62). In our current study, we evaluated mucosal vaccination with DTaP in the coughing rat model of pertussis. To our knowledge, this study is the first to evaluate both IN and OG administered DTaP in the coughing rat model of pertussis. The data presented here suggest that, not only does mucosal immunization protect against bacterial burden, but also against *Bp* induced cough by WBP (**Fig. 2&7**).

Vaccine mediated immunity has been studied in the coughing rat model of pertussis. Utilizing audio tape recorders, Hall *et al* (1998) illustrated that 3 and 5 component aP vaccines administered subcutaneously could protect against *Bp* induced cough (41). wP immunization administered intraperitoneally also decreased the incidence of cough in rats (39). Neither study noted protection against bacterial colonization. Here, in our study, WBP was used to investigate *Bp* induced cough in IN-aP and OG-aP vaccinated rats, as well as measure bacterial burden in the respiratory tract. Our data supports that mucosal administration of DTaP protects against bacterial burden in the respiratory tract and reduced *Bp*-induced cough (**Fig. 2&7**). In addition, IN-aP vaccination reduced bronchial constriction in the lung that is elicited by *Bp* infection (**Fig. 4**). Previous studies evaluating wP and aP vaccines in the rat model of pertussis used one human dose per rat for immunization prior to challenge (39, 41). In an effort to best model an appropriate human to rat dose, we utilized a 1/5^th^ human dose to prime and boost based on the relative sizes of rats compared to mice. We have reported 1/40^th^ human dose as protective in mice and rats are roughly 10x the weight of mice (63). One caveat of this study is that we did not evaluate other human-to-rat titrations of aP. Identification of a minimal protective rat dose would allow for the investigation of vaccine efficacy of new potential antigens/adjuvants in this model (63).

Mucosal immunization has been of particular interest in the pertussis field. We and others have recently evaluated intranasal immunization of DTaP in *Bp* challenged mice (27, 28, 64–66). Intranasally DTaP vaccinated mice were protected against *Bp* challenge and also generated both systemic and mucosal antibodies (27, 28). Live attenuated strain BPZE1 administered intranasally was protective against *Bp* challenge in mice and baboons, and is currently in Phase 2 of clinical studies (33, 34, 67, 68). BPZE1 immunization induces both anti-*Bp* IgG and IgA antibodies systemically and has an increase in resident memory T cells in the lung (33, 69). Oral immunization has also been investigated as a possible vaccination strategy against pertussis. Oral immunization of heat-inactivated *Bp* protected newborns against *Bp* challenge, as well as generated serum and saliva antibody titers (70). Recombinant technologies has led to the development of live attenuated *Salmonella* strains presenting *Bp* antigens (71, 72). Oral immunization of *Salmonella typhimurium aro* vaccine strain harboring the gene for PRN resulted in reduced bacterial colonization in the lung post *Bp* challenge (71). *Salmonella dublin aroA* mutant expressing the gene for FHA was also orally administered as a vaccine in mice (72). Vaccination with this strain induced IgG and IgA antibody titers to FHA in the serum and gut (72). Our current study shows that mucosal immunization not only induced systemic anti-*Bp* IgG but also anti-*Bp* IgA antibodies that likely play role in clearance at the mucosa (**Fig. 4-5, 7**).

CoP is defined as the immune response that is statistically accountable for the observed protection (73). While no CoP has yet to be fully agreed upon against pertussis in humans, anti-PT IgG levels >5 IU/ml are associated with protection in humans (74). In mice, IN administration of DTaP induced anti-*Bp*, anti-FHA, and anti-PT IgG antibodies while wP vaccination induced serum anti-*Bp* IgG antibodies that correlated with protection against *Bp* (28, 57). Here in our study, we generated correlograms between all vaccinated groups to identify correlations between variables in the coughing rat model of pertussis, which has yet to be established (**Fig. 9**). Our results indicate that bacterial clearance in the lung, trachea, and nasal cavity negatively correlate with systemic anti-*Bp* and -PT IgM and anti-*Bp* IgG antibodies in IM-wP and IM-aP immunized rats, while systemic and mucosal antibodies correlated with bacterial clearance in IN-aP and OG-aP vaccinated rats (**Fig. 9**). Antibodies generated following immunization also negatively correlated with a decrease in total cough counts in vaccinated rats (**Fig. 9**). These results suggest that the increase in systemic and mucosal antibodies induced from IN and OG vaccination correlates with protection against *Bp* burden in the respiratory tract and *Bp* induced cough. It is important to note, that OG-aP immunized rats did not generate significant serum antibody titers to the whole bacterium until day 9 post-challenge (**Fig. 4B**). Also, two OG-aP immunized rats had an increase in anti-PT IgG antibody titers in the serum and anti-*Bp* IgA antibodies in the lung (**Fig. 4D, 5A**). We did however detect antigen specific B cells in the bone marrow and 3 of the 4 rats had low levels of IgA in the lung in OG-aP immunized rats (**Fig. S3**). One caveat that should be mentioned is that we did not investigate the T cell responses (Th1/Th2/Th17/Trm/Tem) in rats but future studies will incorporate this into the study design. We have proposed a summary for mechanism for oral vaccination of aP (**Fig. S9**). We hypothesize that this could be because limited amount of vaccine that successfully travels to the gut-associated lymphoid tissue (GALT) for the generation of an immune response. Oral vaccines have to travel through increased pH in the stomach while limited absorption and availability for antigen recognition also occur in the gastrointestinal tract (75). Increase in dose, number of doses, or delivering vaccine in an encased vehicle are potential methods to increase orally vaccinated immune responses. Targeting of vaccine to intestinal M cells for antigen presentation has also been shown to increase oral vaccine efficacy (76). Though we did not study new adjuvants here, we hypothesize adjuvants can aid in stimulating a protective mucosal immune response. These approaches could all potentially increase the efficacy observed through oral vaccination of aP. Furthermore, to deliver the vaccine to the gut, one could envision novel deliver mechanisms such as gelatin coated chewables similar to gummy vitamins that are now popular.

In summary, mucosal vaccination not only protected against bacterial burden in the respiratory tract of challenged rats, but also protected against *Bp* induced cough and respiratory distress measured by WBP (**Fig. 2-3, 7**). It is critical that “next generation pertussis” vaccines protect against bacterial colonization in the lung, nasal cavity, and trachea, as disease manifestations are dependent on bacterial colonization of the lung and trachea, mediated by FHA and fimbriae (77, 78). Both IN and OG immunized rats generated anti-*Bp* specific IgG antibodies in the serum, while IN vaccinated also generated significant anti-*Bp* IgA antibody titers in the nasal cavity following challenge (**Fig. 4-5**). IN-aP and OG-aP immunized rats were protected against acute and total inflammation in the lung (**Fig. 6**). Our data support the potential of a mucosal vaccination against *Bp.*

Further work is needed to fully characterize the immune response generated following IN-aP and OG-aP vaccination in rats, as well as vaccine mediated immunity from vaccination in the coughing rat model of pertussis. T cell immune responses that have been shown to play a role in natural and vaccine mediated immunity against pertussis have yet to be evaluated in the coughing rat model of pertussis due to limited availability of resources to adequately measure T cell responses. Vaccine mediated memory has yet to be evaluated in the coughing rat model of pertussis. Additional research is needed to critically assess vaccine mediated memory, as it is essential that next generation of pertussis vaccines induce longer lasting memory then current vaccines. Future work is also needed to evaluate mucosal immunization against current circulating strains of *Bp* as current strains are genetically divergent from strains of the past, with the goal of making the most efficacious vaccine against *Bp*.

## MATERIALS AND METHODS

### Vaccine composition and administration

INFANRIX (GSK Cat. 58160-810-11) acellular pertussis human vaccine (DTaP) and the National Institute for Biological Standards and Control WHO whole cell pertussis vaccine (NIBSC code 94/532) was used for this study. Vaccines were diluted with endotoxin-free Dulbecco’s PBS (Thermo Fisher Scientific Cat. TMS012A) to a concentration of 1/5^th^ human dose. Vaccines were diluted and administered no more than 1 hr. from composition. The first dose of vaccine was administered to three-week-old (50g) female Sprague-Dawley rats (Charles River Cat. 001CD). At six weeks of age, a booster vaccine of the same dose was administered, followed by *Bp* challenge at eight weeks of age. Intramuscular (IM) vaccinated rats received 100μl in the right thigh muscle of the hind limb. Intranasal (IN) immunized rats were first anesthetized with isoflurane until breathing was minimal. Rats then received 50μl of vaccine in each nostril for a 100μl dose. Oral gavage (OG) vaccinated rats received 100μl dose delivered curved 18 gauge feeding needle (Fisher Scientific Cat. NC9349775). MVC control group received 100μl of the same endotoxin free PBS used to dilute the vaccines in the right thigh muscles of the hind limb. One-week post-prime, two-week post-prime, and one-week post-boost blood was collected via saphenous blood draws for serological analysis. 5mm animal lancets (Fisher Scientific Cat. NC9891620) was used for blood draw. Blood was collected in capillary tubes (Fisher Scientific Cat. NC9059691) for centrifugation. Blood was spun at 15,000x *g* for 3 min., serum collected and stored at −80°C until analysis.

### *Bordetella pertussis* strains and growth conditions

*Bp* strain D420 was cultured on Bordet Gengou (BG) agar (Remel™ Cat. R45232) supplemented with 15% defibrinated sheep blood (Hemostat Laboratories Cat. DSB500) (1). Bacteria cultured BG plates incubated for 48 hrs at 36°C. Using polyester swabs (Puritan Cat. 22-029-574), *Bp* was transferred into 20 ml Stainer-Scholte liquid media (SSM) in new 125 ml flasks (Thermo Fisher Scientific Cat. FB500125) (79). Bacterial cultures were allowed to grow at 36°C for 24 hrs inside a shaking incubator at 180 rpm.

### Intranasal challenge

Vaccinated eight-week-old ~200g female Sprague-Dawley rats were then challenged. *Bp* was grown as illustrated above. Rats were anesthetized with ketamine and xylazine 50-100/5-10 mg/kg and challenged with 10^8^ CFUs in 100μl intranasally, 50μl in each nostril. Body weight of each rat was recorded before bacterial challenge, and body weights were taken post-euthanasia to calculate percent weight change. At days 1 and 9 post challenge, rats were then euthanized. Upon euthanasia blood was collected via cardiac puncture and transferred into ethylenediaminetetraacetic acid (EDTA) (BD Cat. 365974) and serum separation (BD Cat. 026897) tubes. Following cardiac puncture, 250μl of blood was collected into EDTA tubes for flow cytometry and ProCyte (IDEXX) analysis, while remaining blood was collected in serum separation tubes to isolate the serum via centrifugation (15,000x *g* for 3 min) and used for serological and cytokine analysis. To determine bacterial burden in the respiratory tract, the lung and trachea was excised separately and homogenize. Lung weights were recorded following excision before homogenization. Lungs were then collected in gentleMACS C tubes (Miltenyi Biotec Cat. 130-096-334) in 2ml of PBS and homogenized using Miltenyi Biotec tissue dissociator (Cat. 130-095-927). Polytron homogenizer was used to homogenize the trachea in 1 ml PBS. Bacterial burden in the nares was determined by flushing 2mls of sterile 1x PBS through the nares and collected for serial dilution and plating. Serial dilutions of the homogenates and nasal collection were plated on BG plates supplemented with ceftibuten (Sigma-Aldrich Cat. SML0037) 10 μg/ml.

### Serological analysis

Enzyme-linked immunosorbent assays (ELISA) was used to measure antibody titers of vaccinated and infected rats. *Bp* specific whole bacteria ELISA plates were coated with 50 μl of 10^8^ *Bp* grown as mentioned above for infection. Antigen specific antibody titers to PT (List Biological Laboratories #180) were measured by coating ELISA plates with 50 μl of antigen per well. Antigen coated plates incubated over night at 4°C. After incubation, plates were washed with 1x PBS-Tween 20 and blocked with 5% skimmed milk for 2 hrs at 37°C. Following blocking, ELISA plates were washed and serum from the saphenous blood draws and blood collected from cardiac puncture post-euthanasia were serially diluted down the ELISA plate and incubated for 2 hrs at 37°C. To measure respiratory IgA antibody titers in the lung and nasal lavages, lung homogenate supernatant and nasal lavage was added and incubated for 2 hrs at 37°C. After incubation, ELISA plates were washed as described above and 100μl of secondary goat anti-rat IgG (SouthernBiotech Cat. 3030-04), goat anti-rat IgM (SouhternBiotech Cat. 3020-04), or goat anti-rat IgA (MyBioSource Cat. MBS539212) was added to the plates at a dilution of 1:2,000 in PBS + 5% milk and incubated for 1 hr at 37°C. Plates were then washed again and 100 μl *p*-nitrophenyl phosphate substate (Thermo Scientific Cat. 37620) was added and the plate was developed for 30 min at room temperature. After development, colorimetric signal of the ELISA plate at *A*450 was measured by a Biotek Synergy H1 microplate reader. Antibody titers were considered positive if values were higher than the baseline. Baseline value for each sample was set as double the average value of the blank, in which no serum, lung supernatant, or nasal lavage added to these well. Limit of detection was set at 50, and any samples with a titer value less than that were set to 50.

### ELISpot assay

ELISpot assay (ImmunoSpot Cat. mTgG-SCE-1M/2) was used to analyze antigen specific B cells in the bone marrow. The right hind femur of the rat was removed and placed into Dulbecco’s modified Eagle’s medium (DMEM) and frozen at −80°C until analysis. Bones were then thawed in water bath at 37°C, and immediately transferred into spin tubes and spun at 1,000x *g* for 3 min to collect the bone marrow. Bone marrow was passed through a 70μm filter to create a single cell suspension. Cells were centrifuged at 350 x *g* for 5 min and the cell pellet was resuspended in CTL test B Media (ImmunoSpot). D420 was cultured as described above and coated the 96-well ELISpot plate as described by ELISA. The plate incubated overnight at 4°C. Plate was then washed with 1x PBS before cells were added. Three serial dilutions of cells (1.25 × 10^6^, 3.13 × 10^5^, and 1.56 × 10^5^) cells added per well and incubated at 37°C overnight. Rabbit anti-Rat IgG antibody (Abcam Cat. ab6733) was used to replace the anti-murine IgG detection antibody that was with the kit. The rest of the protocol was followed as per the manufacturer’s instructions. ELISpot plates imaged and analyzed using ImmunoSpot S6 Entry analyzer and CTL software.

### Analysis of cough and bronchiole restriction using whole-body plethysmography

Buxco® FinePointe™ Whole Body Plethysmography (WBP) (DSI) was used to quantify respiratory function during infection. Every day following *Bp* challenge and one day before challenge (5:00PM), rat respiratory profiles and coughs were measured. A 5 min acclimation period was used before measuring cough and other respiratory parameters. After acclimation the respiratory profile was recorded for 15 mins for each rat. Coughs were counted and represented over 15 mins. Enhanced pause (PenH) was calculated which represents bronchiole restriction during breathing. Coughs were counted based on box flow changes of the subject with classical cough-like waveforms. Patented fuzzy logic criteria was used to detect and count coughs (80). Each cough in a multi-cough event was counted individually. Frequency (F), Tidal Volume (TVb), Pause (PAU), Minute Volume (MVb), Inspiratory time (Ti), and expiratory time (Te) were also collected and analyzed during the course of infection.

### Histological assessment of the lung

The left lobe of the lung was used for histological assessment. Following excision of the left lobe, the sectioned portion was fixed in 10% formalin 48 hrs at 26 °C. Following fixation, samples were embedded in paraffin and stained with H&E by the WVU Pathology Department. Stained samples were used to characterize and score inflammation of the lung. All scorings were done by a board-certified pathologist (iHisto). Individual scores were based on a standard qualitative scoring criterion: (0 – none, 1 – minimal (rare), 2 – mild (slight), 3 – moderate, 4 – marked, 5 –severe). The presence of neutrophils in the parenchyma, blood vessels, and airway was used to score acute inflammation, and chronic inflammation was characterized by mononuclear infiltrates of the parenchyma, blood vessels, and airway. All examination and scoring were done with no knowledge of the groups.

### ProCyte analysis of blood

Blood from the EDTA tubes was used to analyze white blood cell, neutrophil, and lymphocyte counts. 25-50μl of blood was drawn from the EDTA tubes and analyzed by the Procyte. After ProCyte analysis the rest of the blood was used for flow cytometry analysis.

### Flow cytometry analysis

Blood samples were lysed with 1x Pharmylse buffer (BD Biosciences Cat. 555899) for 20 min at room temperature. Blood samples were vortexed periodically during the 20 min incubation. After lysis, cells were resuspended in RPMI + 10% FBS to neutralize the lysis buffer and centrifuged at 1,000g x 5. Cells were then washed with the RPMI +10% FBS again. Cells were then resuspended in 1%FBS+PBS+5mM EDTA. Blood samples were then blocked with anti-CD32 (BD Pharmingen Cat. 550270) antibody for 15 min at 4°C. After blocking, the cells were labeled; CD45 Alexa flour 700 (Biolegend Cat. 202218), CD161 APC (Biolegend Cat. 205606), CD45R PE Cy7 (eBioscience Cat. 25-0460-82), His48 FITC (eBioscience Cat. 11-0570-82), CD43 PE (Biolegend Cat. 202812), and CD3 VioGreen (Miltenyi Biotec Cat. 130-119-125) (81). Samples were incubated 1 hr at 4°C in the dark. To prepare the lung samples for flow cytometry, the lung homogenate was filtered through a 70 μm cell strainer (BioDesign Cell MicroSives Cat. N70R). The suspension was centrifuged at 1,000 x *g* for 5 min. The pellet was resuspended in Pharmlyse buffer and incubated at 37°C for 2 min. After incubation, the cells were centrifuged at 1,000 x *g* for 5 min, lysis buffer removed, and cells were blocked and labeled with antibody as described for blood samples. Both blood and lung samples were centrifuged at 1,000 x *g* for 5 min and the pellets were resuspended in 0.4% paraformaldehyde and stored overnight at 4°C. Samples were washed with 1x PBS+5mM EDTA+1%FBS and resuspended in 1x PBS+5mM EDTA+1%FBS for analysis. Cell samples were analyzed on a LSR Fortessa and samples were gated and analyzed using FlowJo v10.

### Cytokine analysis

Lung homogenates were centrifuged at 19,000× *g* for 4 min and the resulting supernatant was removed and stored at −80 °C until further analysis. Lung supernatant and serum cytokines were measured using a ProcartaPlex Multiplex Immunoassay kit: Th Complete 14-Plex Rat ProcartaPlex Panel (Thermo Fisher Scientific Cat. EPX140-30120-901) per the manufacturer’s instructions. Cytokines with bead counts less than 35 were invalidated.

### Generation of correlograms

Correlograms were created using R Studio software. Pearson correlation coefficients were calculated between each set of variables listed in the master table and then illustrated in the representative plot for each vaccine route and timepoint.

## Statistical analysis

GraphPad Prizm 7 was used to analyze the data. The minimum biological replicates for the challenge studies were three for MVC control group and four rats per vaccinated groups. For statistical comparisons between vaccinated groups and the MVC control group over the entire course of the infection, two-way analysis of variance (ANOVA) was used with Dunnett’s post hoc test. One-way ANOVA was used for comparison between vaccinated groups and MVC for an individual day or timepoint with Dunnett’s post hoc test. Kruskal-Wallis test with Dunnett’s post hoc test compared between groups for mucosal IgA comparisons. ROUT test was used to identify any potential outliers during cytokine analysis of the lung.

## Data availability

Data requests for figures provided can be addressed to the corresponding author.

## Ethics statement

This challenge study was performed in accordance with our approved protocol by West Virginia University Institutional Animal Care and Use Committee (IACUC) protocol 1811019148.6.

## ACKNOWLEDGMENTS

The work performed in this project was supported by the Vaccine Development Center at WVU-HSC through a Research Challenge Grant no. HEPC.dsr.18.6 from the Division of Science and Research, WV Higher Education Policy Commission. The project was also supported by NIH R01AI137155 (F.H.D). Part of the project was also supported by CDC Contract Broad Agency Announcement (BAA) 75D301-19-R-67835. Flow cytometry experiments were performed in the West Virginia University Flow Cytometry Core Facility, which is supported by the National Institutes of Health equipment grant number S10OD016165 and the Institutional Development Award (IDeA) from the National Institute of General Medical Sciences of the National Institutes of Health under grant numbers P30GM103488 (CoBRE) and P20GM103434 (INBRE).

JMH and GJB performed vaccination and bacterial challenge. JMH and GJB monitored rats by whole body plethysmography. All authors participated in the animal experiments. JMH and TYW prepared flow cytometry samples. JMH and MAW performed cytokine analysis. JMH performed ELISA assays. MAD constructed correlograms. JMH, MB, and FHD contributed to experimental design. JMH wrote manuscript with critical revisions from all authors.

The authors would also like to thank Jacqueline Karakiozis, Brice Hickey, and Mary Tomago-Chesney (Pathology/Histology Core Facility) for the preparation of lung slides and staining lung slides with H&E for scoring analysis. The authors would also like to thank Dr. Kathleen Brundage (WVU Flow Cytometry & Single Cell Core Facility) for assisting in flow cytometry and equipment instruction.

The authors would also like to acknowledge that figures S1 and S8 were created with BioRender.com

## References

1. Bordet J, Gengou O. 1906. Le Microbe de la Coqueluche. Les Ann l’Institut Pasteur 20:731–741.

2. Mattoo S, Cherry JD. 2005. Molecular pathogenesis, epidemiology, and clinical manifestations of respiratory infections due to *Bordetella pertussis* and other *Bordetella* subspecies. Clin Microbiol Rev 18:326–382.

3. Melvin JA, Scheller E V., Miller JF, Cotter PA. 2014. *Bordetella pertussis* pathogenesis: Current and future challenges. Nat Rev Microbiol2014/03/13. 12:274–288.

4. Kapil P, Merkel TJ. 2019. Pertussis vaccines and protective immunity. Curr Opin Immunol 59:72–78.

5. CDC, Ncird. Immunology and Vaccine-Preventable Diseases – Pink Book – Pertussis.

6. Pittman M. 1991. History of the development of pertussis vaccine. Dev Biol Stand1991/01/01. 73:13–29.

7. Hill Elam-Evans LD, Yankey D, Singleton JA, Kang Y. HA. 2017. Vaccination Coverage Among Children Aged 19–35 Months — United States, 2017. MMWR Morb Mortal Wkly Rep 2018 1123–1128.

8. Yeung KHT, Duclos P, Nelson EAS, Hutubessy RCW. 2017. An update of the global burden of pertussis in children younger than 5 years: a modelling study. Lancet Infect Dis 17:974–980.

9. Klein NP, Bartlett J, Fireman B, Baxter R. 2016. Waning Tdap Effectiveness in Adolescents. Pediatrics 137:e20153326–e20153326.

10. Klein NP, Bartlett J, Rowhani-Rahbar A, Fireman B, Baxter R. 2012. Waning Protection after Fifth Dose of Acellular Pertussis Vaccine in Children. N Engl J Med 367:1012–1019.

11. Klein NP, Bartlett J, Fireman B, Rowhani-Rahbar A, Baxter R. 2013. Comparative Effectiveness of Acellular Versus Whole-Cell Pertussis Vaccines in Teenagers. Pediatrics 131:e1716–e1722.

12. Althouse BM, Scarpino S V. 2015. Asymptomatic transmission and the resurgence of *Bordetella pertussis*. BMC Med 13:146.

13. Templeton KE, Scheltinga SA, van der Zee A, Diederen BMW, van Kruijssen AM, Goossens H, Kuijper E, Claas ECJ. 2003. Evaluation of real-time PCR for detection of and discrimination between *Bordetella pertussis*, *Bordetella parapertussis*, and *Bordetella holmesii* for clinical diagnosis. J Clin Microbiol 41:4121–6.

14. Warfel JM, Zimmerman LI, Merkel TJ. 2014. Acellular pertussis vaccines protect against disease but fail to prevent infection and transmission ina nonhuman primate model. Proc Natl Acad Sci U S A 111:787–792.

15. Wendelboe AM, Van Rie A, Salmaso S, Englund JA. 2005. Duration of Immunity Against Pertussis After Natural Infection or Vaccination. Pediatr Infect Dis J 24:S58–S61.

16. Wilk MM, Misiak A, McManus RM, Allen AC, Lynch MA, Mills KHG. 2017. Lung CD4 Tissue-Resident Memory T Cells Mediate Adaptive Immunity Induced by Previous Infection of Mice with *Bordetella pertussis*. J Immunol 199:233–243.

17. Chen K, Magri G, Grasset EK, Cerutti A. 2020. Rethinking mucosal antibody responses: IgM, IgG and IgD join IgA. Nat Rev Immunol. Nature Research.

18. Goodman YE, Wort AJ, Jackson FL. 1981. Enzyme-linked immunosorbent assay for detection of pertussis immunoglobulin A in nasopharyngeal secretions as an indicator of recent infection. J Clin Microbiol 13:286–292.

19. Tuomanen EI, Zapiain LA, Galvan P, Hewlett EL. 1984. Characterization of antibody inhibiting adherence of *Bordetella pertussis* to human respiratory epithelial cells. J Clin Microbiol 20:167.

20. Hellwig SMM, Van Spriel AB, Schellekens JFP, Mooi FR, Van de Winkel JGJ. 2001. Immunoglobulin A-mediated protection against *Bordetella pertussis* infection. Infect Immun 69:4846–4850.

21. Thomas MG, Redhead K, Lambert HP. 1989. Human Serum Antibody Responses to *Bordetella pertussis* Infection and Pertussis Vaccination. J Infect Dis 159:211–218.

22. Marcellini V, Piano Mortari E, Fedele G, Gesualdo F, Pandolfi E, Midulla F, Leone P, Stefanelli P, Tozzi AE, Carsetti R. 2017. Protection against Pertussis in Humans Correlates to Elevated Serum Antibodies and Memory B Cells. Front Immunol 8:1158.

23. Mills KH, Barnard A, Watkins J, Redhead K. 1993. Cell-mediated immunity to *Bordetella pertussis*: role of Th1 cells in bacterial clearance in a murine respiratory infection model. Infect Immun 61:399–410.

24. Dunne A, Ross PJ, Pospisilova E, Masin J, Meaney A, Sutton CE, Iwakura Y, Tschopp J, Sebo P, Mills KHG. 2010. Inflammasome Activation by Adenylate Cyclase Toxin Directs Th17 Responses and Protection against *Bordetella pertussis*. J Immunol 185:1711–1719.

25. Warfel JM, Merkel TJ. 2013. *Bordetella pertussis* infection induces a mucosal IL-17 response and long-lived Th17 and Th1 immune memory cells in nonhuman primates. Mucosal Immunol 6:787–796.

26. Solans L, Locht C. 2019. The role of mucosal immunity in pertussis. Front Immunol. Frontiers Media S.A.

27. Boehm DT, Wolf MA, Hall JM, Wong TY, Sen-Kilic E, Basinger HD, Dziadowicz SA, Gutierrez M de la P, Blackwood CB, Bradford SD, Begley KA, Witt WT, Varney ME, Barbier M, Damron FH. 2019. Intranasal acellular pertussis vaccine provides mucosal immunity and protects mice from *Bordetella pertussis*. npj Vaccines 4.

28. Wolf MA, Boehm DT, DeJong MA, Wong TY, Sen-Kilic E, Hall JM, Blackwood CB, Weaver KL, Kelly CO, Kisamore CA, Bitzer GJ, Bevere JR, Barbier M, Damron FH. 2020. Intranasal immunization with acellular pertussis vaccines results in long-term immunity to *Bordetella pertussis* in mice. Infect Immun IAI.00607-20.

29. Maurer H, Höfler K, Hilbe W, Huber E. 1979. Preliminary findings with oral whooping cough vaccination in young infants. Wien Hlin Wochenschr.

30. Baumann E, Binder BR, Falk W, Huber EG, Kurz R, Rosanelli K. 1985. Development and clinical use of an oral heat-inactivated whole cell pertussis vaccine. Dev Biol Stand 61:511–6.

31. Guzman CA, Brownlie RM, Kadurugamuwa J, Walker MJ, Timmis KN. 1991. Antibody responses in the lungs of mice following oral immunization with *Salmonella typhimurium aroA* and invasive *Escherichia coli* strains expressing the filamentous hemagglutinin of *Bordetella pertussis*. Infect Immun 59:4391–4397.

32. Lim A, Ng JKW, Locht C, Alonso S. 2014. Protective role of adenylate cyclase in the context of a live pertussis vaccine candidate. Microbes Infect 16:51–60.

33. Skerry CM, Mahon BP. 2011. A live, attenuated *Bordetella pertussis* vaccine provides long-term protection against virulent challenge in a murine model. Clin Vaccine Immunol2010/12/08. 18:187–193.

34. Thorstensson R, Trollfors B, Al-Tawil N, Jahnmatz M, Bergström J, Ljungman M, Törner A, Wehlin L, Van Broekhoven A, Bosman F, Debrie AS, Mielcarek N, Locht C. 2014. A phase I clinical study of a live attenuated *Bordetella pertussis* vaccine - BPZE1; a single centre, double-blind, placebo-controlled, dose-escalating study of BPZE1 given intranasally to healthy adult male volunteers. PLoS One 9:e83449.

35. Lin A, Apostolovic D, Jahnmatz M, Liang F, Ols S, Tecleab T, Wu C, van Hage M, Solovay K, Rubin K, Locht C, Thorstensson R, Thalen M, Loré K. 2020. Live attenuated pertussis vaccine BPZE1 induces a broad antibody response in humans. J Clin Invest 130:2332–2346.

36. Hornibrook JW, Ashburn LL. 1939. A Study of Experimental Pertussis in the Young Rat. Public Heal Reports 54:439.

37. Woods DE, Franklin R, Cryz SJ, Ganss M, Peppler M, Ewanowich C. 1989. Development of a rat model for respiratory infection with *Bordetella pertussis*. Infect Immun 57:1018–1024.

38. Hall E, Parton R, Wardlaw AC. 1994. Cough production, leucocytosis and serology of rats infected intrabronchially with *Bordetella pertussis*. J Med Microbiol 40:205–213.

39. Parton R, Hall E, Wardlaw AC. 1994. Responses to *Bordetella pertussis* mutant strains and to vaccination in the coughing rat model of pertussis. J Med Microbiol 40:307–312.

40. Hall E, Parton R, Wardlaw AC. 1997. Differences in coughing and other responses to intrabronchial infection with *Bordetella pertussis* among strains of rats. Infect Immun 65:4711–4717.

41. Hall E, Parton R, Wardlaw AC. 1998. Responses to acellular pertussis vaccines and component antigens in a coughing-rat model of pertussis. Vaccine 16:1595–603.

42. Hall E, Parton R, Wardlaw AC. 1999. Time-course of infection and responses in a coughing rat model of pertussis. J Med Microbiol 48:95–98.

43. Hall JM, Kang J, Kenney SM, Wong TY, Bitzer GJ, Kelly CO, Kisamore CA, Boehm DT, DeJong MA, Allison M, Sen-Kilic E, Horspool AM, Bevere JR, Barbier M, Heath Damron F. 2021. Re-investigating the coughing rat model of pertussis to understand *Bordetella pertussis* pathogenesis. bioRxiv 2021.04.02.438291.

44. Ross PJ, Sutton CE, Higgins S, Allen AC, Walsh K, Misiak A, Lavelle EC, McLoughlin RM, Mills KHG. 2013. Relative Contribution of Th1 and Th17 Cells in Adaptive Immunity to *Bordetella pertussis*: Towards the Rational Design of an Improved Acellular Pertussis Vaccine. PLoS Pathog 9:e1003264.

45. Ryan M, Murphy G, Ryan E, Nilsson L, Shackley F, Gothefors L, Øymar K, Miller E, Storsaeter J, Mills KH. 1998. Distinct T‐cell subtypes induced with whole cell and acellular pertussis vaccines in children. Immunology 93:1–10.

46. Ausiello CM, Urbani F, la Sala A, Lande R, Cassone A. 1997. Vaccine- and antigen-dependent type 1 and type 2 cytokine induction after primary vaccination of infants with whole-cell or acellular pertussis vaccines. Infect Immun 65:2168–2174.

47. Mahon BP, Sheahan BJ, Griffin F, Murphy G, Mills KH. 1997. Atypical disease after *Bordetella pertussis* respiratory infection of mice with targeted disruptions of interferon-gamma receptor or immunoglobulin mu chain genes. J Exp Med 186:1843–51.

48. Barbic J, Leef MF, Burns DL, Shahin RD. 1997. Role of gamma interferon in natural clearance of *Bordetella pertussis* infection. Infect Immun 65:4904–4908.

49. Mills KHG, Ryan M, Ryan E, Mahon BP. 1998. A murine model in which protection correlates with pertussis vaccine efficacy in children reveals complementary roles for humoral and cell-mediated immunity in protection against *Bordetella pertussis*. Infect Immun 66:594–602.

50. Redhead K, Watkins J, Barnard A, Mills KHG. 1993. Effective immunization against *Bordetella pertussis* respiratory infection in mice is dependent on induction of cell-mediated immunity. Infect Immun 61:3190–3198.

51. Sawal M, Cohen M, Irazuzta JE, Kumar R, Kirton C, Brundler M-A, Evans CA, Wilson JA, Raffeeq P, Azaz A, Rotta AT, Vora A, Vohra A, Abboud P, Mirkin LD, Cooper M, Dishop MK, Graf JM, Petros A, Klonin H. 2009. Fulminant pertussis: A multi-center study with new insights into the clinico-pathological mechanisms. Pediatr Pulmonol 44:970–980.

52. Paddock CD, Sanden GN, Cherry JD, Gal AA, Langston C, Tatti KM, Wu K, Goldsmith CS, Greer PW, Montague JL, Eliason MT, Holman RC, Guarner J, Shieh W, Zaki SR. 2008. Pathology and Pathogenesis of Fatal *Bordetella pertussis* Infection in Infants. Clin Infect Dis 47:328–338.

53. Morse SI, Morse JH. 1976. Isolation and properties of the leukocytosis- and lymphocytosis-promoting factor of *Bordetella pertussis*. J Exp Med 143:1483–1502.

54. Morse SI. 1965. STUDIES ON THE LYMPHOCYTOSIS INDUCED IN MICE BY *BORDETELLA PERTUSSIS*. J Exp Med 121:49–68.

55. Morse SI, Riester SK. 1967. Studies on the leukocytosis and lymphocytosis induced by *Bordetella pertussis*. I. Radioautographic analysis of the circulating cells in mice undergoing pertussis-induced hyperleukocytosis. J Exp Med 125:401–8.

56. Heininger U, Klich K, Stehr K, Cherry JD. 1997. Clinical findings in *Bordetella pertussis* infections: results of a prospective multicenter surveillance study. Pediatrics 100:E10.

57. Blackwood CB, Sen-Kilic E, Boehm DT, Hall JM, Varney ME, Wong TY, Bradford SD, Bevere JR, Witt WT, Damron FH, Barbier M. 2020. Innate and adaptive immune responses against *Bordetella pertussis* and *Pseudomonas aeruginosa* in a murine model of mucosal vaccination against respiratory infection. Vaccines 8:1–21.

58. Correlogram. https://www.r-graph-gallery.com/correlogram.html

59. Bewick V, Cheek L, Ball J. 2003. Statistics review 7: Correlation and regression. Crit Care. BioMed Central.

60. Althouse BM, Scarpino S V. 2015. Asymptomatic transmission and the resurgence of *Bordetella pertussis* https://doi.org/10.1186/s12916-015-0382-8.

61. Klein NP, Bartlett J, Fireman B, Aukes L, Buck PO, Krishnarajah G, Baxter R. 2017. Waning protection following 5 doses of a 3-component diphtheria, tetanus, and acellular pertussis vaccine. Vaccine 35:3395–3400.

62. Nakamura K, Shinoda N, Hiramatsu Y, Ohnishi S, Kamitani S, Ogura Y, Hayashi T, Horiguchi Y. 2019. BspR/BtrA, an Anti-σ Factor, Regulates the Ability of *Bordetella bronchiseptica* To Cause Cough in Rats. mSphere https://doi.org/10.1128/msphere.00093-19.

63. Boehm DT, Hall JM, Wong TY, DiVenere A, Sen-Kilic E, Bevere JR, Bradford SD, Blackwood CB, Elkins C, DeRoos KA, Gray MC, Cooper CG, Varney ME, Maynard JA, Hewlett EL, Barbier M, Damron FH. 2018. Evaluation of adenylate cyclase toxoid antigen in acellular pertussis vaccines using a *Bordetella pertussis* challenge model in mice. Infect Immun IAI.00857-17.

64. Shi W, Kou Y, Jiang H, Gao F, Kong W, Su W, Xu F, Jiang C. 2018. Novel intranasal pertussis vaccine based on bacterium-like particles as a mucosal adjuvant. Immunol Lett 198:26–32.

65. Allen AC, Wilk MM, Misiak A, Borkner L, Murphy D, Mills KHG. 2018. Sustained protective immunity against *Bordetella pertussis* nasal colonization by intranasal immunization with a vaccine-adjuvant combination that induces IL-17-secreting T RM cells. Mucosal Immunol 11:1763–1776.

66. Ryan EJ, Mcneela E, Murphy GA, Stewart H, O’Hagan D, Pizza M, Rappuoli R, Mills KHG. 1999. Mutants of *Escherichia coli* heat-labile toxin act as effective mucosal adjuvants for nasal delivery of an acellular pertussis vaccine: Differential effects of the nontoxic AB complex and enzyme activity on Th1 and Th2 cells. Infect Immun 67:6270–6280.

67. Locht C, Papin JF, Lecher S, Debrie A-S, Thalen M, Solovay K, Rubin K, Mielcarek N. 2017. Live Attenuated Pertussis Vaccine BPZE1 Protects Baboons Against *Bordetella pertussis* Disease and Infection. J Infect Dis2017/05/23. 216:117–124.

68. Solans L, Debrie A-S, Borkner L, Aguiló N, Thiriard A, Coutte L, Uranga S, Trottein F, Martín C, Mills KHG, Locht C. 2018. IL-17-dependent SIgA-mediated protection against nasal *Bordetella pertussis* infection by live attenuated BPZE1 vaccine. Mucosal Immunol 11:1753–1762.

69. Lin A, Apostolovic D, Jahnmatz M, Liang F, Ols S, Tecleab T, Wu C, van Hage M, Solovay K, Rubin K, Locht C, Thorstensson R, Thalen M, Loré K. 2020. Live attenuated pertussis vaccine BPZE1 induces a broad antibody response in humans. J Clin Invest 130.

70. Baumann E, Binder BR, Falk W. 1985. Development and clinical use of an oral heat-inactivated whole cell pertussis vaccine. Dev Biol Stand VOL. 61:511–516.

71. Strugnell R, Dougan G, Chatfield S, Charles I, Fairweather N, Tite J, Li JL, Beesley J, Roberts M. 1992. Characterization of a *Salmonella typhimurium* aro vaccine strain expressing the P.69 antigen of *Bordetella pertussis*. Infect Immun 60.

72. Molina NC, Parker CD. 1990. Murine antibody response to oral infection with live aroA recombinant *Salmonella dublin* vaccine strains expressing filamentous hemagglutinin antigen from *Bordetella pertussis*. Infect Immun 58.

73. Plotkin SA. 2010. Correlates of protection induced by vaccination. Clin Vaccine Immunol. American Society for Microbiology.

74. Murphy TV, Slade BA, Broder KR, Kretsinger K, Tiwari T, Joyce PM I, JK, Brown K MJAC on IP, Prevention. C for DC and. 2008. Prevention of Pertussis, Tetanus, and Diphtheria Among Pregnant and Postpartum Women and their Infants Recommendations of the Advisory Committee on Immunization Practices (ACIP). MMWR Recomm Reports 1–51.

75. Frizzell H, Woodrow KA. 2020. Biomaterial Approaches for Understanding and Overcoming Immunological Barriers to Effective Oral Vaccinations. Adv Funct Mater 30:1907170.

76. Azizi A, Kumar A, Diaz-Mitoma F, Mestecky J. 2010. Enhancing oral vaccine potency by targeting intestinal M cells. PLoS Pathog. Public Library of Science.

77. Kilgore PE, Salim AM, Zervos MJ, Schmitt H-J. 2016. Pertussis: Microbiology, Disease, Treatment, and Prevention. Clin Microbiol Rev 29:449–86.

78. Scheller E V., Cotter PA. 2015. *Bordetella* filamentous hemagglutinin and fimbriae: critical adhesins with unrealized vaccine potential. Pathog Dis 73:ftv079.

79. Stainer DW, Scholte MJ. 1970. A Simple Chemically Defined Medium for the Production of Phase I *Bordetella pertussis*. J Gen Microbiol1970/10/01. 63:211–220.

80. Lomask J, Larson R. 2006. United States Patent USOO7104962B2. US 7,104,962 B2.

81. Barnett-Vanes A, Sharrock A, Birrell MA, Rankin S. 2016. A Single 9-Colour Flow Cytometric Method to Characterise Major Leukocyte Populations in the Rat: Validation in a Model of LPS-Induced Pulmonary Inflammation https://doi.org/10.1371/journal.pone.0142520.

